# Mapping the temporal and functional landscape of Sonic Hedgehog signaling reveals new insights into early human forebrain development

**DOI:** 10.1101/2025.05.20.654466

**Authors:** Veranika Panasenkava, Jules Garreau, Hélène Guyodo, Farah Diab, Ariane Mahieux, Wilfrid Carré, Yann Verres, Emmanuelle Jullion, Sylvie Odent, Christèle Dubourg, Paul Rollier, Lauryane Dubos, Vanessa Ribes, Anne Camus, Marie de Tayrac, Valérie Dupé

## Abstract

The early patterning of the anterior neuroectoderm constitutes a fundamental blueprint for human brain development, orchestrated by multiple signaling pathways. Among them, Sonic Hedgehog (SHH) plays a key influence. However, the transcriptional programs it engages remain poorly defined due to the limited accessibility of human brain tissue. To address this, we established a human induced pluripotent stem cells-derived model of early forebrain differentiation, enabling a precise dissection of SHH-driven transcriptional programs over time. RNA sequencing revealed dynamic transcriptomic landscapes governing forebrain neuroectoderm specification and dorsoventral patterning. In addition, pharmacological perturbation of SHH signaling allowed to identify an extended collection of novel forebrain regionalization markers, including several previously unrecognized dorsal and ventral determinants. By combining *in vivo* human datasets with functional mouse studies, we enhanced the biological relevance of this extended network of putative SHH-regulated genes and long non-coding RNAs in shaping early forebrain architecture. This work advances our understanding of the temporal dynamics of SHH signaling in human neurodevelopment and provide critical molecular insights into midline brain malformations. It offers a promising foundation for advancing molecular diagnosis of complex rare genetic disorders.

**GRAPHICAL ABSTRACT:** 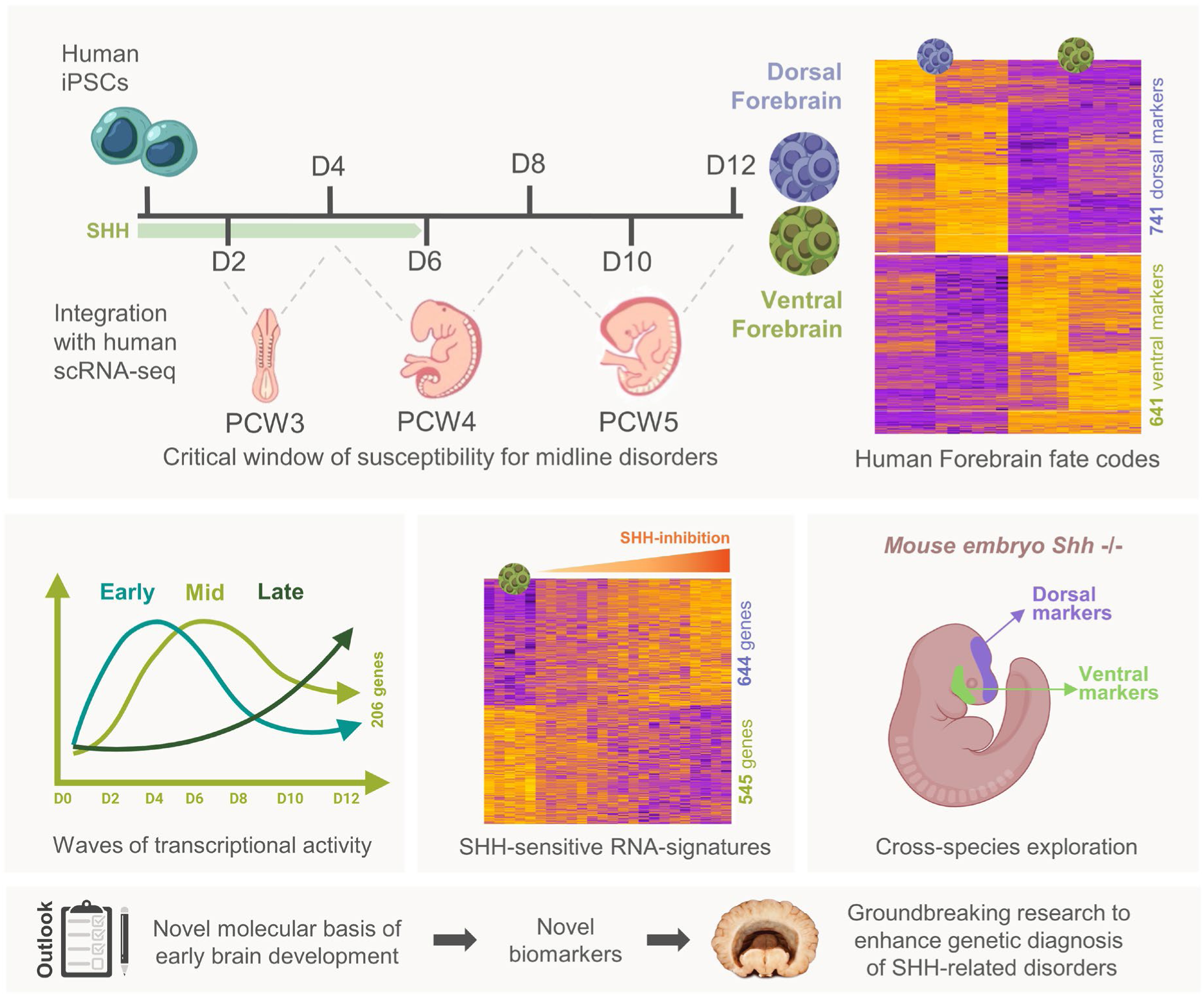

## INTRODUCTION

Early brain development begins with neuroectoderm regionalization into distinct compartments, a process orchestrated by a combination of spatial and temporal molecular cues. The anterior neural plate gives rise to the forebrain (prosencephalon), which splits into the telencephalon (future cortex) and diencephalon (future hypothalamus and pituitary). These domains are defined through intricate signaling interactions that define the molecular identity and future differentiation of neural progenitors (Puelles, 2024; Deutsch Guerrero et al., 2025). A key regulator in this early patterning is the morphogen Sonic Hedgehog (SHH), which is initially secreted by the prechordal plate (PCP) to promote ventral identity in the overlying neuroectoderm (Wilson & Houart, 2004).

Subsequently, SHH expression becomes localized at the ventral midline of the neural plate, establishing a gradient the highest concentration in the floor plate, which regulates the activity of Gli transcriptions factors to control cell fate specification (Bai et al., 2004; Kohtz et al., 1998, Fuccillo et al., 2004). These, in turn, initiate region-specific transcriptional programs essential for ventral forebrain development, beginning with the activation of NKX2.1 (Douceau et al., 2023; Bertrand & Dahmane, 2006; Shimamura et al., 1995; Ratié et al., 2013; Ware et al., 2014). Conversely, dorsal territories are marked by the expression of other transcription factors as PAX6 (Fuccilo et al., 2006), highlighting the tightly regulated spatial dichotomy imposed by SHH Signaling.

This transcriptional regulation is highly sensitive to perturbations in SHH activity (Mercier et al., 2013; Placzek and Briscoe, al., 2018). In mice, loss of *Shh* results in the complete absence of ventral forebrain structures (Chiang, 1996; Ishibashi et al., 2002), while moderate reductions lead to midline malformations such as pituitary defects (Guyodo et al., 2024). Moreover, subtle change in the timing, intensity, or duration of SHH signaling can trigger graded malformations of the rostral brain as early as E8.25 (Geng et al., 2008; Zhang et al., 2006; Christ et al., 2012). In human, comparable disruptions during the third to fifth post-conception weeks (PCW) are though to underlie a spectrum of congenital brain disorders ranging from pituitary stalk interruption syndrome to holoprosencephaly (Wang et al., 2017; Mercier et al. 2011). Variants in *SHH* itself, or in genes involved in its activity, are among the most frequent causes of these disorders (Dubourg et al., 2018; Lavillaureix et al., 2024, Bando et al., 2023).

Despite substantial knowledge derived from animal models, the molecular logic by which SHH instructs early ventral identity in the human anterior neuroectoderm remains largely elusive. While *in vitro* studies have described SHH function in human forebrain neural progenitors, (Hunt et al., 2023; Tchieu et al., 2017; Xiang et al., 2017; Birey et al., 2017; Cederquist et al., 2019), it remains unclear how SHH signaling precisely regulate the earlier transcriptional programs. Moreover, the extent to which this morphogenic control influences cell fate decisions underlying congenital midline brain defects is still not fully understood determined.

To address this gap, we developed an optimized 2D differentiation system from hiPSCs to generate anterior neuroectoderm-like cells (AN), exhibiting either dorsal (dAN) or ventral (vAN) characteristics. Using transcriptomic profiling combined with graded pharmacological perturbation of SHH signaling, we systematically characterized the gene expression landscape along the dorsal–ventral axis. This approach allowed us to dissect how spatial identity emerges across the human anterior neuroectoderm. In parallel, we validated the biological relevance of several novel candidate genes by analyzing their expression and function during early brain development in mouse embryos.

## RESULTS

### An efficient protocol for neuroectodermal specification and dorsal–ventral patterning from human induced pluripotent stem cells

To study the molecular mechanism of anterior neuroectodermal (AN) development in a reproducible and time-controlled way (Figure 1A), we established a 12-day 2D monolayer differentiation protocol with human induced pluripotent stem cells (hiPSCs) for two independent wild-type lines LON (Gaignerie et al., 2018) and WTC (Conklin Lab, San Franciso) (Supplementary Figure 1A, B). Neuroectoderm induction was achieved through dual SMAD inhibition (dSMADi) with LDN-193189 (LDN) and SB431542 (SB), inhibiting BMP and TGF-β pathways respectively, throughout the 12 days of culture (Tchieu et al., 2017). To promote anterior identity, the WNT inhibitor, XAV939 (XAV) was applied from day 0 to day 5 (Qi et al., 2017; Major et al., 2017; Alekseenko et al., 2022).

**Figure 1:**
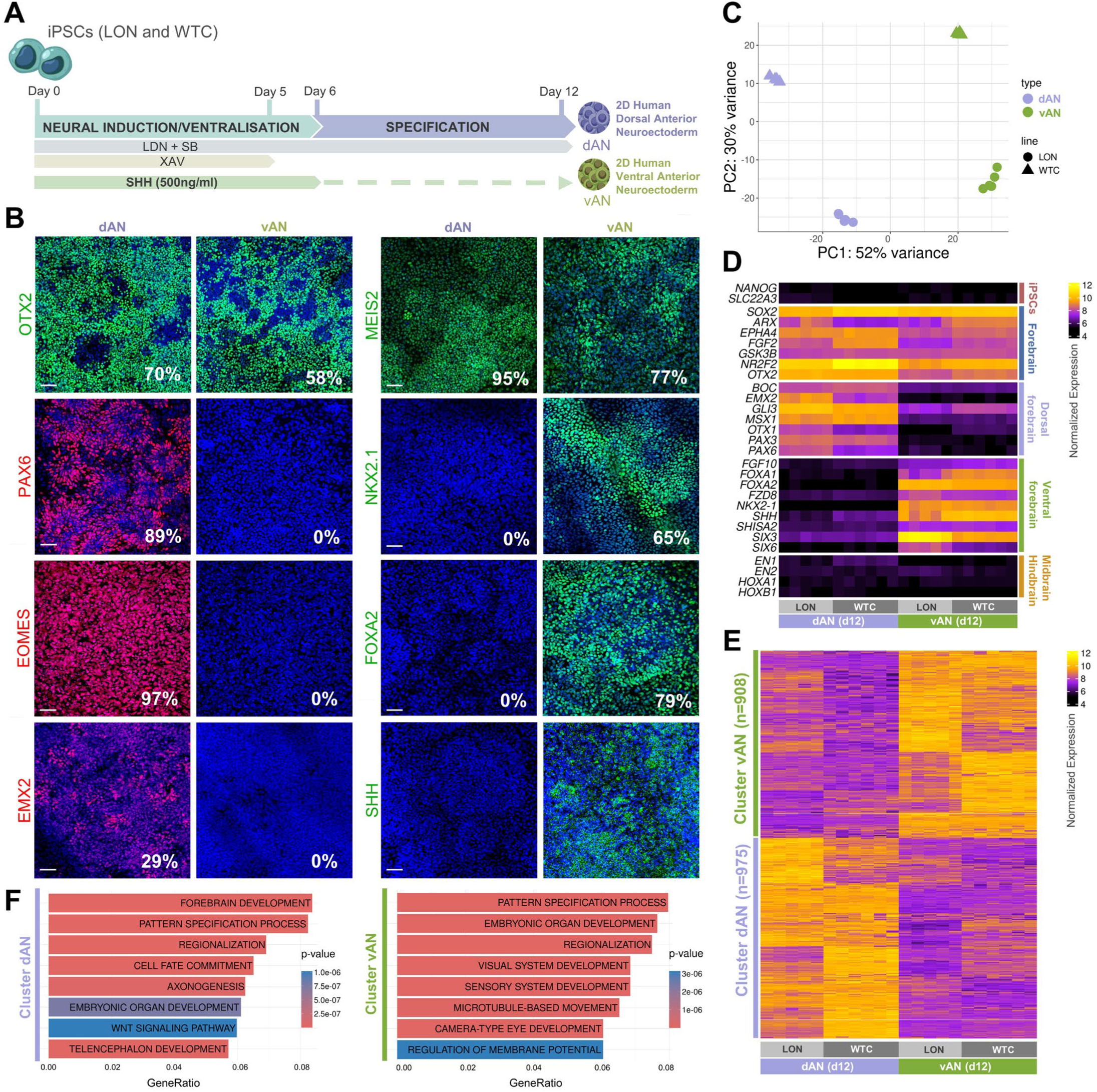
Efficient generation of dorsal and ventral anterior neuroectoderm from human iPSCs. (A) Schematic illustration of the differentiation protocol for generating dorsal (dAN) and ventral (vAN) anterior neuroectoderm from hiPSCs over a 12-day period. Neural induction and ventralization are initiated by a combination of small molecules: LDN193189 (500 nM), SB431542 (10 µM), from day 0 to day 12, and XAV939 (5 µM) from day 0 to day 5. Additional exposure to human recombinant Sonic Hedgehog (SHH, 500 ng/ml) from day 0 to day 6 selectively drives ventral patterning in vAN cultures. (B) Representative Immunofluorescent images at day 12 show distinct marker expression in dAN and vAN cells. Markers analyzed include OTX2 (anterior neuroectoderm marker), PAX6 and EMX2 (dorsal forebrain markers), EOMES (early neural cortical marker), MEIS2 (forebrain marker), NKX2.1 (ventral forebrain marker), FOXA2 (floor plate marker), and SHH (ventral marker). Expression levels are quantified as the percentage of positively stained cells relative to DAPI (mean values from 5 independent images). Scale bar: 40 µm. (C) Principal Component Analysis (PCA) of day 12 dAN and vAN samples (LON and WTC lines) from 5 replicates revealed clear segregation by identity and lineage. PC1 (52% total variance) strongly distinguished dAN versus vAN conditions (one-way ANOVA p.value of 8.9e-12), while PC2 (30% total variance) separates samples by cell lineage origin (one-way ANOVA p.value of 3.1e-12). (D) Heatmap displaying normalized RNA-seq expression profiles of day 12 dAN and vAN samples. Samples were manually ordered by type (vAN/dAN) and cell line (LON/WTC). Marker genes sets specific for hiPSCs, forebrain, ventral/dorsal Forebrain, Midbrain and Hindbrain identity were curated from literature. (E) Heatmap of scaled normalized RNA-seq data showing differentially expressed genes (DEGs) between day 12 dAN and vAN samples (|log2FC| **≥** 1, FDR < 0.01), across 2 hiPSC lines (LON, WTC). Both genes and samples were hierarchical clustered (cluster vAN and cluster dAN), two cluster were identified and then delimited using cutree R function with k=2. (F) Brain development-related Gene Ontology terms among the top 20 most enriched (in term of gene ratio) biological process for both DEGs clusters (dAN and vAN). Color scale indicates gene sets enrichment statistical significance.

We first characterized the cell identities of differentiated LON cells at day 12 using immunostaining for forebrain markers (Figure 1B, Supplementary Figure S1C). By this stage, cells had formed small neural rosettes and displayed forebrain identity, with the majority expressing the pan-forebrain markers OTX2 (70%) and MEIS2 (95%) (dAN in Figure 1B and Supplementary figure 1C) (Leung et al., 2022). Most cells also expressed dorsal forebrain markers PAX6 (89%) and EOMES (97%), supporting their classification as dorsal anterior neuroectoderm (dAN) (Figure 1B, Supplementary Figure 1C). Interestingly, 29% of dAN cells were positive for EMAX2, EMX2, a dorsal marker specific to the telencephalon, indicating partial regionalization within the dorsal forebrain lineage (Figure 1B, Supplementary Figure 1C). As expected, these dAN did not expressed ventral markers NKX2-1, FOXA2 or SHH (Figure 1B).

To generate ventral AN-like cells (vAN), we initiated SHH signaling from day 0, as early SHH activation is critical for ventral forebrain specification (Danjo et al., 2011; Krajka et al., 2021). To sustain this activation, cells were continuously exposed to recombinant human SHH (500 ng/ml) from day 0 to day 6. By day 12, the pan-forebrain markers OTX2 (58%) and MEIS2 (77%) were expressed, along with the ventral forebrain marker NKX2-1 (65%), a key indicator of hypothalamic fate, and the floor plate marker FOXA2 (79%), both of which are direct targets of SHH signaling. Dorsal markers (PAX6, TBR2, EMX2) were undetectable in vAN cells. Notably, high level of extracellular SHH protein was observed at day 12 (vAN in Figure 1B, Supplementary Figure 1D). Together, these expression profiles of vAN confirm the successful establishment of predominant hypothalamic characteristic cells, the structure that emerges from the ventral forebrain.

Reverse-transcription quantitative PCR (RT-qPCR) assays confirmed consistent dorsal (*PAX6, SOX1, TBR1*) and ventral (*SHH, NKX2-1, FOXA1*) gene expressions patterns in dAN and vAN cells respectively across the two iPSC lines (Supplementary figure 1E). Importantly, it showed that vAN cells autonomously produced SHH by day 12, even though SHH treatment ended by day 6, indicating effective endogenous activation of the SHH pathway by the recombinant SHH protein.

We then performed bulk RNA-seq analysis at day 12 under both dAN and vAN conditions for the two iPSC lines, LON and WTC. Principal Component Analysis (PCA) of the top 3,000 most variable genes identified the main source of variability (Figure 1C). PC1 (52%) primarily separated cells based on dAN and vAN conditions (one-way ANOVA, p=8.85×10^-12^), while PC2 (30%) reflected differences between cell lines (one-way ANOVA, p=3.07×10^-12^). This indicates that the primary source of variability arises from the difference between dAN and vAN conditions.

We assessed the expression of brain-specific markers by curating gene lists from the literature (Figure 1D) (Leung et al., 2022). Markers associated with the AN region, such as *ARX, EPHA4, FGF2, GSK3B, NR2F2* and *OTX2*, were expressed in both dAN and vAN cells. Dorsal markers such as *BOC, EMX2, GLI3, MSX1, OTX1, PAX3 and PAX6* showed high expression in dAN cells. Conversely, the high expression of ventral markers such as *FGF10, FOXA1, FOXA2, FZD8, NKX2-1, SHH, SHISA2, SIX3 and SIX6* in vAN cells confirms their acquisition of ventral identity, aligning with the activation of the SHH signaling pathway. Notably, the transcription factors specific to the midbrain (*EN1, EN2)* and hindbrain (*HOXA1, HOXB1*) were not found to be expressed under dAN or vAN conditions.

We next performed a differential expression analysis comparing vAN and dAN samples (Figure 1E, Supplementary table 1), which identified 1,883 differentially expressed genes (DEGs) (FDR < 0.01, |log2FC| >= 1). Of these, 975 were highly expressed in dAN samples and showed low expression in vAN (cluster dAN), while 908 genes exhibited the opposite pattern, low expression in dAN and high expression in vAN (cluster vAN). GeneOntology enrichment analysis indicates that both gene sets are associated with early forebrain regionalization process (Forebrain development, Pattern specification, Regionalization, Cell fate commitment; Figure 1F, Supplementary table 2A-B). Specifically, cluster dAN is strongly linked with telencephalic development, whereas cluster vAN is enriched for gene involved in eye development and primary cilia function. Notably, many of these genes have not previously been described as playing a function in anterior neuroectoderm development. However, for several of them, such as *ZFHX4* (cluster dAN), and *TRIM9*, *EPHB1*, and the lncRNA, *SFTA3* (cluster vAN), we have characterized their expression patterns in the mouse embryo during early forebrain development. A feedback loop within *SFTA3-Nkx2.1* gene duplex is essential for buffering *Nkx2.1* expression to maintain lung development in mouse (Herriges et al., 2017). A similar function may be implicated in forebrain development as we found that *SFTA3* shows strong spatial correlation with *NKX2.1 mRNA* in mouse forebrain development (Supplementary Figure 1F).

Overall, these gene expression analyses validate the protocol’s effectiveness to reliably generate ventral and dorsal neuroectoderm cell populations with forebrain identity across two genetic backgrounds, underscoring the robustness of the differentiation strategy. Importantly, this study establishes distinct transcriptomic profiles for vAN and dAN conditions, revealing several novel markers specific to the dorsal or ventral forebrain.

### A two-phase transcriptional program governing forebrain specification revealed by time-resolved transcriptomics

In order to grasp the dynamics transcriptional changes underlying differentiation toward vAN and dAN fates, we performed a time-course RNA-seq analysis. Samples were collected every other day from day 2 to day 12, including two biological replicates for each of the two iPSC lines (LON and WTC) (Figure 2A). PCA of the 3,000 most variable genes showed strong clustering of samples based on experimental conditions rather than cell line, indicting high reproducibility across replicates. The first principal component (PC1), which explained 36% of the total variance, aligned with the temporal progression of differentiation (one-way ANOVA, p-value = 1.2e-23). Furthermore, as early as day 2, samples began to diverge along the PC2, capturing 23% of the variance (one-way ANOVA, p-value = 1.9e-15), and clearly separated according to dAN and vAN culture conditions (Figure 2B).

**Figure 2:**
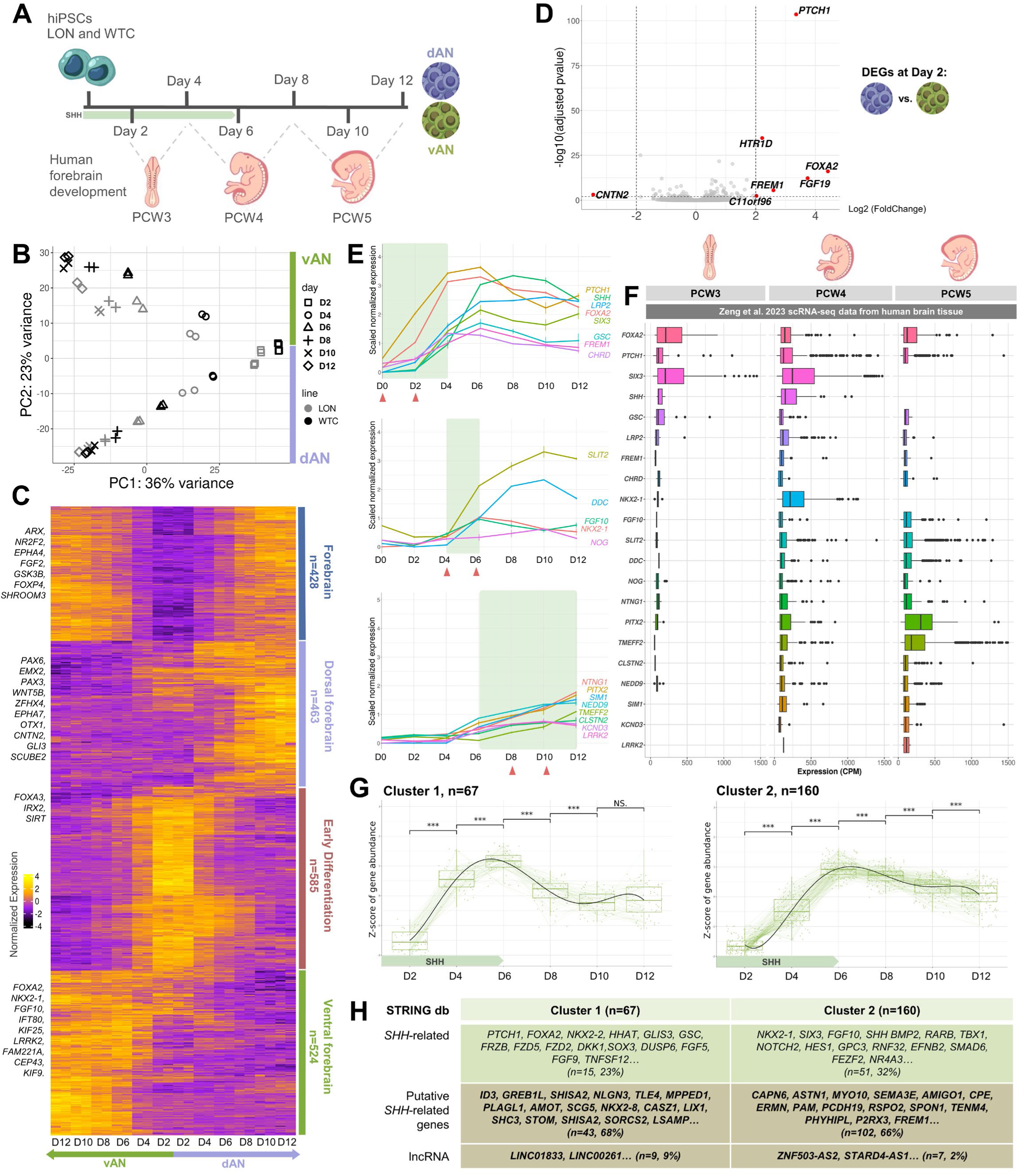
Stage-specific characterization of hiPSC-derived dorsal and ventral anterior neuroectoderm differentiation. (A) Schematic timeline of the *in vitro* protocol (dAN and vAN) aligned with key stages of human embryonic development. Days 2-4: Carnegie Stages (CS) 8–9 (∼post-conception weeks [PCW] 3); days 4-8: CS 10–11 (PCW 4–5); days 8-12: CS 12-13 (PCW 5–6). (B) PCA of RNA-seq time course (day 2, 4, 6, 8, 10, 12) of dAN and vAN differentiation from two hiPSC lines (LON, WTC; n=5 replicates/line). PCA reveals: PC1 (36% total variance), temporal progression (p=1.2×10⁻²³, ANOVA); PC2 (23% total variance) dorsal-ventral divergence (p.value of 1.9e-15, ANOVA). (C) Heatmap of scaled RNA-seq expression (WTC line) for the top 1000 genes correlated with either PC1 and/or PC2 (Pearson correlation to PCA eigenvectors). Genes were hierarchically clustered, four cluster were identified and then delimited using cutree R function with k=4. Samples were manually ordered by type (dAN/vAN) and timepoint (days 2-12). Gene clusters were annotated as: forebrain (n=428), ventral forebrain (n=524), dorsal forebrain (n=463) and early differentiation markers (n=585). (D) Volcano plot of dAN versus vAN DEGs at day 2. Labeled genes meet thresholds of |log2FC| ≥ 2 and FDR < 0.01. Dashed lines indicate significant cutoffs. (E) Temporal expression patterns of curated marker genes in WTC-derived cultures. Values are mean of scaled normalized expression values among timepoints. Genes are grouped by activation timing with respectively from top to bottom: early (pre-day 4), intermediate (days 4-6), and late (post-day 6). (F) Expression levels (counts per million - CPM) of 2E gene markers in human forebrain and neuronal cells (PCW 3-5), as annotated in the original single-cell study (Zeng et al., 2023). (G) Temporal expression patterns of DEGs in WTC samples. Cluster 1 (n=67) and 2 (n=160) represent distinct DEG groups with significant expression changes across at least 2 consecutive timepoints on WTC samples. Each plot displays: individual gene trajectories with Z-score normalized expression (thin lines); Cluster trendline: Mean expression pattern (bold line); Statistical significance: Wilcoxon signed-rank test p-value (“***”=0.001, “**”=0.01, “*”=0.05, “NS.”=Non-significant) between time points. (H) Genes from Cluster 1 and Cluster 2 (2G) were analyzed using the STRING database to identify known or predicted relationships with SHH pathway (SHH-related), and to highlight novel genes and lncRNAs.

Next, we selected the top 2,000 genes most strongly correlated. With the PC1 and PC2 eigenvectors, based on Pearson correlation across all samples. These gens were visualized in a heatmap (Figure 2C for WTC; Supplementary Figure 2A for LON; Supplementary Table 3A-B). Unsupervised clustering identified four distinct expression clusters, each exhibiting a characteristic temporal profile aligned with successive neuroectodermal states and/or lineage commitments stages.

The first cluster, labeled *“Early Differentiation”*, included 585 genes that reached peak expression at the onset of differentiation, followed by a gradual downregulation from day 4 to day 12 in both dAN and vAN conditions (Figure 2C). This initial transcriptional wave was marked by genes like *FOXA3*, a key player in anterior neural fate induction prior to gastrulation (Seiliez et al., 2006), *IRX2*, involved in early forebrain patterning (Rodríguez-Seguel et al., 2009), and *SIRT1*, which promotes neural survival in early stages (Cai et al., 2015). GO enrichment revealed significant associations with translational processes, consistent with increased protein synthesis requirements during early developmental transitions (Supplementary Figure 2B, Supplementary Table 4A).

In contrast, the *“Forebrain”* cluster consisted of 428 genes that remained transcriptionally silent at early time points and became progressively upregulated from day 4 onward under both dAN and vAN conditions. This transcriptional wave included key regulators of anterior neuroectodermal identity, such as *ARX*, *NR2F2*, *EPHA4*, *FGF2*, and *GSK3B*. Key examples include *FOXP4*, essential for forebrain formation and neural progenitor specification, and *SHROOM3*, involved in shaping neuroepithelial architecture (Plageman et al., 2011). These results point to the initiation of anterior neuroectodermal specification from day 4 in both lineages (Figure 2; Supplementary Figure 2B; Supplementary Table 4B).

The *“Dorsal Forebrain”* cluster, encompassing 463 genes, showed marked specifically upregulation in the dAN lineage from day 4 onward. This cluster was enriched in canonical dorsal forebrain markers, including *PAX6*, *EMX2*, *PAX3*, *WNT5B*, *ZFHX4*, *EPHA7*, *OTX1*, and *CNTN2*. Additionally, it included SHH pathway antagonists such as *GLI3* and *SCUBE2*. GO term analysis underscored significant associations with forebrain development and neural progenitor identity, supporting the establishment of dorsal fate in dAN cells (Supplementary Figure 2B, Supplementary Table 4C).

In contrast, the *“Ventral Forebrain”* cluster contained 524 genes that were specifically upregulated in the vAN lineage. This cluster included early ventral markers such as *FOXA2*, *NKX2-1*, and *FGF10*, alongside a notable enrichment in genes related to primary cilia, including *IFT80*, *KIF25*, *LRRK2*, *FAM221A*, *CEP43*, and *KIF9*. GO term analysis highlighted a strong signature of cilium assembly and function, consistent with active SHH signal transduction. Additionally, several components of the WNT signaling pathway were also enriched in this cluster, indicating convergence of multiple developmental signals in ventral patterning (Supplementary Figure 2B, Supplementary Table 4D).

Collectively, these findings outline a two-phase model of early forebrain differentiation. During the initial phase (days 1 to 4), cells exhibit high proliferative activity and a progressive loss of pluripotency. By day 4, both dAN and vAN populations have commited to an anterior neuroectodermal lineage. Following this commitment, the lineages begint: vAN cells progressively adopt ventral identity, while dAN cells initiate dorsal programs. By day 6, many canonical markers for dorsal and ventral regionalization are already expressed, marking a critical transition in forebrain patterning.

### Sequential SHH-Driven Gene Expression Waves Map the Differentiation Trajectory of Forebrain Patterning

To assess the functional significance of SHH-induced transcriptional dynamics during ventral anterior neuroectoderm (vAN) differentiation, we conducted a time-resolved analysis of DEGs in both WTC (Figure 2) and LON (Supplementary Figure S2) iPSC-derived models. DEGs were systematically identified across the 12-day time course, with early (Day 2) expression changes visualized by volcano plots (Figure 2D) and longitudinal trajectories analyzed using the degPatterns function from DEGReport (Figure 2E, G; Supplementary Figure S2C). To contextualize these transcriptional dynamics within developmental context, we compared our dataset to: (i) human embryonic single-cell RNA-seq (PCW3–5; Zeng et al., 2023) (Figure 2F); (ii) curated mouse forebrain expression profiles; and (iii) a SHH interactome derived from the STRING database (confidence ≥ 0.4) (Figure 2H).

Although SHH was present in the culture medium from day 0, the early transcriptional response was limited. Volcano plot analysis at day 2 (Figure 2D; Supplementary Table 5) identified only six significantly upregulated genes in vAN cells (FDR < 0.01, |log₂FC| ≥ 2). Among these were canonical SHH targets, *PTCH1*, *FOXA2*, and *FGF19*, whose expression in the mouse aligns with embryonic day 8.0 (E8.0) (Ishibashi & McMahon, 2002; Vincentz et al., 2005). Notably, *FREM1*, a known SHH-responsive gene in neural crest cells (McLaughlin et al., 2023), was also upregulated at this early stage. This indicates that SHH signaling, despite limited transcriptional breadth, had already begun to activate early targeted programs.

By day 4, a broader and more coordinated transcriptional response to SHH became evident. Of the nine dynamic profiles identified, “*Profile 1”* captured a sharp and transient transcriptional surge of a group of 67 genes. They were upregulated between day 4 and day 6, followed by a downregulation phase by day 8 (FDR < 0.01, |log₂FC| ≥ 2; Supplementary Tables 5, 6A). This cluster included *SHH* itself, indicating a potential shift toward autocrine signaling. It also contained *SIX3*, a known positive regulator of *SHH* expression (Carlin et al., 2012), and *LRP2*, a co-receptor involved in SHH uptake and response in the floor plate (Christ et al., 2012). Additional ventral forebrain regulators such as *GSC* and *CHRD* - genes known to contribute to early forebrain fate acquisition - were also enriched in this group (Lewis & Tam, 2006; Anderson et al., 2002). These dynamics are consistent with a transcriptional shift occurring around mouse E8.25, marking the onset of early neural patterning.

STRINGdb based analysis of *“Profile 1*” identified 15 genes (25%) with established SHH interactions, including *NKX2-2* and *GSC*, corroborating the module’s biological relevance. Among the 45 genes not previously linked to SHH signaling, several, such as *ID3*, *GREB1L*, *SHISA2*, and *NLGN3,* exhibit documented roles in neural progenitor cell fate. Notably, this cluster included six long non-coding RNAs (lncRNAs), two of which - *LINC00261 and LINC01833* - emerged has putative SHH signaling components. *LINC00261*, which is positionally conserved near *FOXA2*, likely contributes to neuroectoderm specification through a mechanism similar to that described during endoderm formation, where it promotes *FOXA2* expression via SMAD2/3 recruitment (Jiang et al., 2015, PMID: 25843708; Swarr et al., 2019). The function of *LINC01833*, positioned near *SIX3*, remains poorly defined – though its genomic suggests a potential regulatory role.

*“Profile 2”* revealed a cohort of 71 genes that were upregulated by day 4 and maintained high expression levels throughout subsequent differentiation. This sustained wave included *NKX2-1*, *FGF10*, *DDC*, *NOG*, and *SLIT2*, all of which are involved in ventral identity specification, proliferation, and early neuronal differentiation. Functional network analysis showed that 32% of Profile 2 genes are linked to SHH signaling, either directly or through first-degree interactors. Notably, several genes in this group - *RGMA*, *SPON1*, *CAPN6-* are associated with neuroepithelial morphogenesis or patterning, yet their roles in forebrain formation remain unexplored. Altogether, this profile reflects a consolidation phase of ventral identity, aligning with embryonic stages E9.0–E9.5 (Figure 2g, H).

*“Profile 3”* encompassed ∼300 genes that showed a progressively upregulated expression from day 6 to day 12 (FDR < 0.01, |log₂FC| ≥ 1; Supplementary Tables 6C-E). This gene set included *NTNG1*, *SIM1*, *PITX2*, *TMEFF2*, *NEDD9*, *KCND3*, *LRRK2* and *CLSTN,* all of which are associated with neurogenesis, neuronal migration, cytoskeletal remodeling, and early circuit formation. This profile aligns temporally with mouse developmental stages E10.0 and beyond, when progenitor zones begin generating distinct neuronal subtypes (Figure 2E).

To further validate the temporal significance of these gene sets, we compared them to human embryonic scRNA-seq data. Profile 1 genes (e.g. *FOXA2*, *GSC*, *PTCH1*) aligned with transcriptional programs active at PCW3, consistent with early forebrain patterning; Profile 2 genes (e.g. *NKX2-1*, *FGF10*, *SLIT2*, *DDC*) overlapped with the emergence of medial ganglionic eminence (MGE) and ventral diencephalic signatures at PCW4; and Profile 3 genes became most prominent at PCW5, corresponding to neural lineage commitment and early circuit specification (Figure 2F).

In summary, our data reveal a sequential cascade of SHH-responsive transcriptional programs driving vAN differentiation, transitioning from early ligand responsiveness to the establishment of regional identity and neuronal fate. The enrichment of known SHH targets and emergence of novel candidates, including non-coding RNAs, underscores the complexity and precision of SHH-regulated gene networks during early human neurodevelopment.

### Graded Cyclopamine Inhibition Demonstrates SHH’s Morphogen Action in Ventral Forebrain Patterning

We investigated how SHH signaling shapes human ventral forebrain fate by treating vAN cultures with increasing concentrations of Cyclopamine (CyP), a well-established antagonist of the SHH receptor Smoothened. Building on previous work on animal model showing that CyP induces a range of forebrain defects proportional to the degree of SHH pathway inhibition (Mercier et al., 2013), we administrated CyP at graded doses (0.125µM, 0.25µM, 0.5µM, and 1µM) from day 6 to day 12, after endogenous SHH induction (Figure 3A). Changes in cellular identity were assessed by double immunostaining for PAX6 and NKX2-1 (Figure 3B). The results revealed a striking dose-dependent shift in cell fate as CyP concentration increased: the proportion of NKX2-1-positive cells progressively declined, while PAX6-positive cells increased. In untreated vAN cultures, 65% of cells expressed NKX2-1, with no detectable PAX6. The lowest CyP dose (0.125µM) led to the emergence of a small PAX6-positive population (6%) and a reduction in NKX2-1-positive cells to 32%. This trend intensified with higher CyP concentrations: at 0.25µM, PAX6 rose to 20% and NKX2-1 fell to 15%; at 0.5µM, PAX6 reached 47% and NKX2-1 was reduced to 13%; and at 1µM, PAX6 peaked at 57%, while NKX2-1 dropped to 8% (Figure 3B-C). No further changes were observed beyond this concentration, indicating a saturation effect (data not shown).

**Figure 3:**
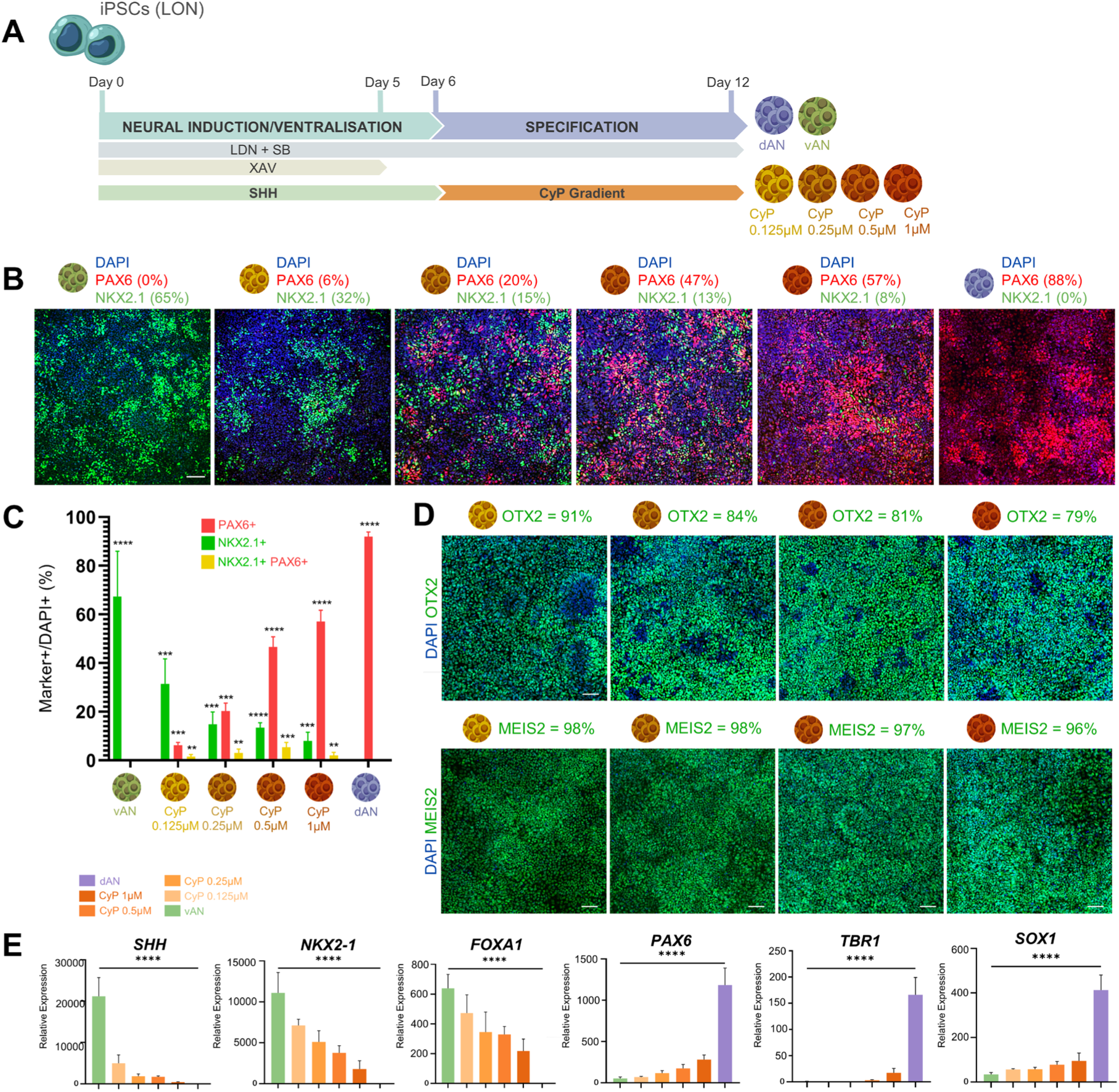
Cyclopamine dose-dependent inhibition of SHH signaling during hiPSC-derived ventral anterior neuroectoderm culture. (A) Schematic representation of SHH pathway modulation in vAN cultures using different Cyclopamine (CyP) concentrations (1 µM to 0.125 µM) from day 6. (B) Representative confocal images showing immunofluorescent staining of PAX6 and NKX2.1 under increasing CyP concentrations (1 µM to 0.125 µM), vAN and dAN conditions. Progressive CyP treatments shifts cellular identity from ventral (High NKX2.1) to dorsal (high PAX6). Scale bar: 40 µm. Data represent the percentage of DAPI-positive cells co-expressing PAX6 or NKX2.1, calculated as mean from five replicates per condition. (C) Analysis of PAX6 and NKX2.1-positive cells at day 12 across vAN, dAN, and CyP gradient (1 µM to 0.125 µM) conditions. Data show the percentage of DAPI-positive cells that express either PAX6 or NKX2.1 or both, averaged across five replicates for each condition. Statistical significance is indicated with **p < 0.01, ***p < 0.001, ****p < 0.0001 (one-way ANOVA). (D) Confocal immunofluorescent images of OTX2 and MEIS2 neuroectodermal markers across CyP concentrations (0.125 µM to 1 µM). Positively stained cells (%) relative to DAPI expression levels are quantified as the percentage of positively stained cells relative to DAPI (mean values from 5 images/condition). Scale bar: 40 µm. (E) Quantitative analysis of dorsoventral patterning under SHH modulation of LON-derived vAN cultures. RT-qPCR profiling of dorsal (*PAX6, TBR1 and SOX1*) and ventral (*SHH*, *NKX2.1*, and *FOXA1*) across vAN, dAN, and CyP gradient (1 µM to 0.125 µM) conditions. Data show normalized relative expression levels, with significance indicated as ****p < 0.0001 calculated from five replicates per condition (one-way ANOVA).

These findings demonstrate that SHH pathway inhibition elicits a robust, dose-dependent shift from ventral to dorsal neuroectodermal identity. Notably, only a small subset of NKX2-1-positive cells co-expressed PAX6 (Figure 3C), supporting that SHH enforces a binary fate decision, driving cells toward either a ventral or dorsal trajectory, rather than promoting intermediate state. Importantly, expression of anterior neuroectodermal markers *OTX2* and *MEIS2* remained unaffected by SHH inhibition (Figure 3D), indicating that SHH specifically governs dorsal–ventral identity, but not anterior, patterning in this context.

These immunostaining results were further supported by qPCR analysis performed on both iPSC lines (Figure 3E; Supplementary Figure 4), which revealed a dose-dependent reduction in the expression of SHH target genes, including *SHH*, *NKX2-1*, and *FOXA1*. Conversely, the expression of dorsal markers such as *PAX6*, *TBR1*, and *SOX1* increased progressively with higher CyP concentrations.

In conclusion, our findings demonstrate that the vAN model responds to SHH inhibition in a dose-dependent manner, marked by a progressive downregulation of ventral transcription factors and a corresponding upregulation of dorsal markers. This confirms SHH as key morphogenetic regulator of dorsal–ventral patterning in the developing human forebrain and highlights the value of this model in accurately capturing the molecular consequences of altered SHH signaling.

### Cyclopamine treatments uncovers SHH dosage-sensitive transcriptomic signatures

To investigate how varying SHH inhibition shapes transcriptional identity, we performed RNA-seq analysis at day 12 of differentiation, comparing ventral untreated ventral anterior neuroectoderm (vAN) cultures with those vAN exposed to increasing doses of CyP (Figure 4). PCA of the 3,000 most variable genes revealed a clear segregation between untreated and CyP-treated samples along PC1 (42% of variance; one-way ANOVA, *p* = 8.95e–8), highlighting distinct molecular profiles (Figure 4A). Notably, CyP-treated samples exhibited a dose-dependent distribution along PC1 (*p* = 7.73e–8), with lower CyP concentrations clustering closer to vAN group and higher doses inducing a progressive shift in transcriptomic signatures. Each cyclopamine dose produced a distinct transcriptomic signature, reflecting the molecular state of the cells according to the level of SHH pathway inhibition. These observations indicate a robust, graded molecular response to SHH inhibition, reflecting dynamic changes in cell state as SHH pathway activity decreases.

**Figure 4:**
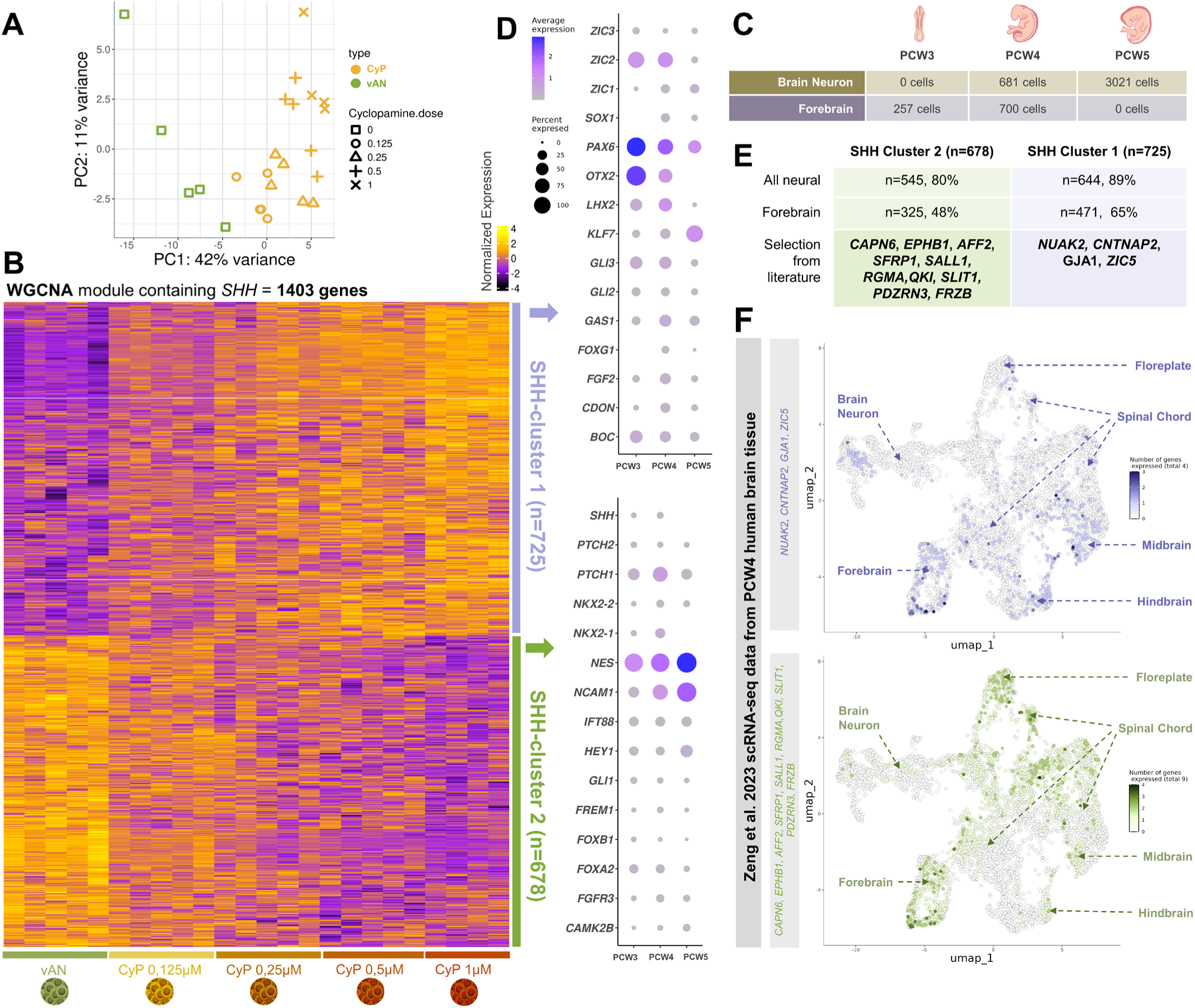
Gene Co-expression networks reveal SHH-dependent modules sensitive to CyP. (A) PCA analysis of RNA-seq data from LON-vAN samples at day 12, treated with graded doses of CyP (0 to 1 µM). PC1 (42% of total variance) strongly correlates with Cyclopamine dosage (one-way ANOVA p.value of 7.7e-8), demonstrating dose-dependent transcriptional shifts. n = 5 replicates per dose of CyP. (B) Heatmap showing scaled normalized expression of the genes with proper HGNC symbol within the WGCNA module containing *SHH* (1,403 genes). Hierarchical clustering was applied to genes, while sample where manually organized based on Cyclopamine dosage. Two cluster were identified and then delimited using cutree R function with k=2. (C) Number of cells in (Zeng et al., 2023) single-cell data annotated by the authors as forebrain and brain neuron at PCW 3, 4, and 5. (D) Average expression and percentage of cells expressing key genes from SHH-cluster 1 and SHH-cluster 2 in human scRNA-seq data at PCW 3, 4, and 5, across forebrain and brain neuron cell types. Color scale represent genes average expression while dot size represent the percentage of cells expressing the genes (no dot equal to 0%). (E) Genes from SHH-clusters 1 (n = 725) and 2 (n = 678) were cross-referenced with human forebrain scRNA-seq data at PCW4 (Zeng et al., 2023), revealing that 89% (n = 644) and 80% (n = 545), respectively, are expressed during early human forebrain development. Selected genes from SHH-cluster 1 (*NUAK2, CNTNAP2, GJA1, ZIC5) and* SHH*-*cluster 2 (*CAPN6, EPHB1, AFF2, SFRP1, SALL1, RGMA, QKI, SLIT1, PDZRN3, FRZB*) were manually curated from the literature based on their potential and previously uncharacterized roles in forebrain development and SHH-related pathways. (F) UMAPs of human central nervous system cells at PCW4 (Zeng et al., 2023) scRNA-seq data annotated by the authors. Cells are colored by the number of genes expressed among SHH-cluster 1-related genes (*NUAK2, CNTNAP2, GJA1, ZIC5*; violet) and SHH-cluster 2-related genes (*CAPN6, EPHB1, AFF2, SFRP1, SALL1, RGMA, QKI, SLIT1, PDZRN3*; green).

To identify gene networks underlying this graded response, we conducted weighted gene co-expression network analysis (WGCNA), which identified a module of 1,656 highly co-expressed, CyP-sensitive genes - including *SHH* itself (Figure 4B; Supplementary Table 9A). After filtering for HGNC-approved gene symbols, the final module comprised 1,403 genes.

Hierarchical clustering further subdivided this module into two distinct subgroups, SHH-cluster 1 and SHH-cluster 2 - each displaying opposing expression dynamics. SHH-cluster 1 (678 genes) mirrored the expression pattern of *SHH* and known targets such as *NKX2-1* and *PTCH1*, indicating co-regulation by SHH signaling. GO enrichment analysis revealed involvement in biological processes including pattern specification, regionalization, cilium organization, and neurogenesis regulation (Supplementary Table 10A). Notably, this SHH-cluster 1 also included the lncRNAs *LINC01833* and *LINC00261*, previously identified as enriched in vAN compared to dAN conditions, further supporting their SHH-dependence.

In contrast, SHH-cluster 2 (725 genes) showed an inverse expression pattern, with progressive upregulation correlating with increasing CyP concentrations. This cluster was enriched for dorsal forebrain markers, including *PAX6*, *EMX2*, *GAS1*, *GLI2*, *GLI3*, *OTX2*, *SOX1*, and *BOC,* consistent with a shift toward dorsal identity under SHH inhibition. GO analysis confirmed their roles in pattern specification, forebrain differentiation, and neurogenesis (Supplementary Table 10B).

Collectively, these results reveal a broader repertoire of genes - encompassing both previously characterized and novel coding and non-coding transcripts – that are responsive to SHH dosage. This expands our understanding of the transcriptional networks that shape early anterior neuroectodermal development and underscores the value of SHH gradient manipulation as a powerful approach for generating distinct transcriptomic signatures and dissecting morphogen-driven patterning.

### Previously uncharacterized SHH-responsive genes implicated in Human anterior neuroectoderm and forebrain patterning

Within the two SHH-responsive clusters, many genes had not previously been linked to early neuroectoderm development. To evaluate their biological relevance, we examined their *in vivo* expression using the publicly available scRNA-seq dataset from the developing human brain (Zeng et al., 2023; Figure 4C). As an initial validation, we selected 15 known ventral neuroectoderm markers from SHH-Cluster 1 and 15 dorsal markers from SHH-Cluster 2 (Figure 4D). Their expression was assessed across PCW3 (257 cells), 4 (1,381 cells), and 5 (3,021 cells), focusing on cells annotated as forebrain. All selected markers were detected in forebrain cells, with robust expression observed by PCW4. Notably, *SHH* and *NKX2-1* were expressed only at PCW3 and PCW4, underscoring their early and transient roles during human forebrain development. These findings support the validity of this dataset for exploring *in vivo* gene expression dynamics during early neurodevelopment.

To expand our investigation, we assessed the expression of all genes from SHH- Clusters 1 and 2 at PCW4, within the neural cell population (n = 6,298). This included forebrain (700 cells), brain neurons (681), midbrain (393), hindbrain (511), spinal cord (3,768), and spinal motor neurons (245), visualized using UMAP embeddings. Remarkably, 89% of SHH-Cluste 1 genes and 80% of SHH-Cluster 2 genes were expressed in at least one neural subpopulation. In the forebrain compartment, 65% of SHH-Cluster 1 and 48% of SHH-Cluster 2 genes were detected (Figure 4E–F; Supplementary Figure 5), supporting, in vi vivo, the biological relevance of our *in vitro*- derived SHH-sensitive clusters.

Based on an extensive literature review, we curated a panel of 14 genes that either (i) had not been previously reported as expressed in the anterior neuroectoderm, or (ii) lacked any known functional link to SHH signaling during forebrain development. This panel included 4 genes from SHH-Cluster 1 (*CNTNAP2*, *GJA1*, *NUAK2*, *ZIC5*) and 10 genes from SHH-Cluster 1 (*CAPN6*, *SALL1*, *SPON1*, *SFRP1*, *QKI*, *PDZRN3*, *FRZB*, *RGMA*, *AFF2*, *SLIT1*) (Figure 4E; Supplementary Figure 5).

We then analyzed the spatial expression of these genes using UMAP projections of human embryonic scRNA-seq data at PCW4 (Figure 4F). Genes in Cluster 1 displayed regional expression, with high levels in the forebrain and detectable presence in the midbrain, and hindbrain, when comparing with the anterior–dorsal (*EMX2*) markers, they were co-expressed. In contrast, SHH-Cluster 2 genes exhibited strong localized expression predominantly in the forebrain and floorplate, with minimal expression in the spinal cord. We next compared the spatial expression patterns of these 10 candidate genes with that of the canonical anterior–ventral (*NKX2-1*). Notably, SHH- Cluster 1 genes showed strong co-localization with *NKX2-1* (Supplementary Figure 5). Among these genes, none have been previously linked to early forebrain development or SHH signaling — except *FRZB* and *SFRP1*, which have been linked to SHH activity during hindbrain development (Liu et al., 2017).

Overall, this systematic comparison between newly identified SHH-sensitive genes and *in vivo* human scRNA-seq data further validates the developmental relevance of our SHH-responsive gene clusters. It reveals a set of previously uncharacterized transcripts likely involved in anterior neuroectoderm patterning. These findings not only strengthen the significance of our *in vitro* differentiation model but also identify promising candidate regulators for future functional studies of early human forebrain development.

### ***In vivo*** functional validation of Newly Identified SHH Target Genes using a mouse model

To complement our *in-silico* analysis, we examined the spatial expression and SHH dependency of these 14 newly identified genes during early brain development using the Shh⁻/⁻ mouse model (Hamdi-Rozé et al., 2020). In situ hybridization was performed on wild-type and mutant embryos with custom RNA probes targeting *Capn6, Sall1, Spon1, Sfrp1, Qki, Rgma, Pdzrn3, Frzb, Aff2, Slit1, Cntnap2, Gja1, Nuak2, and Zic5*.

By embryonic day 9.0 (E9.0), both the telencephalon and diencephalon are properly specified in *Shh^-/-^* embryos; however, these regions exhibit distinct morphological differences compared to Wild-type (WT) embryos, reflecting impaired forebrain development (Chiang et al., 1996; Ishibashi, et al., 2002). At this stage, the forebrain is typically divided into 2 regions, anteriorly, is the telencephalon, and posteriorly, the presumptive diencephalon, which appears as a narrow region flanked by two constrictions that separate it from the telencephalon and midbrain. Notably, the ventral constriction between the diencephalon and midbrain is markedly deeper in *Shh^-/-^* embryos compared to their WT counterparts (black arrowhead in Figure 5).

**Figure 5:**
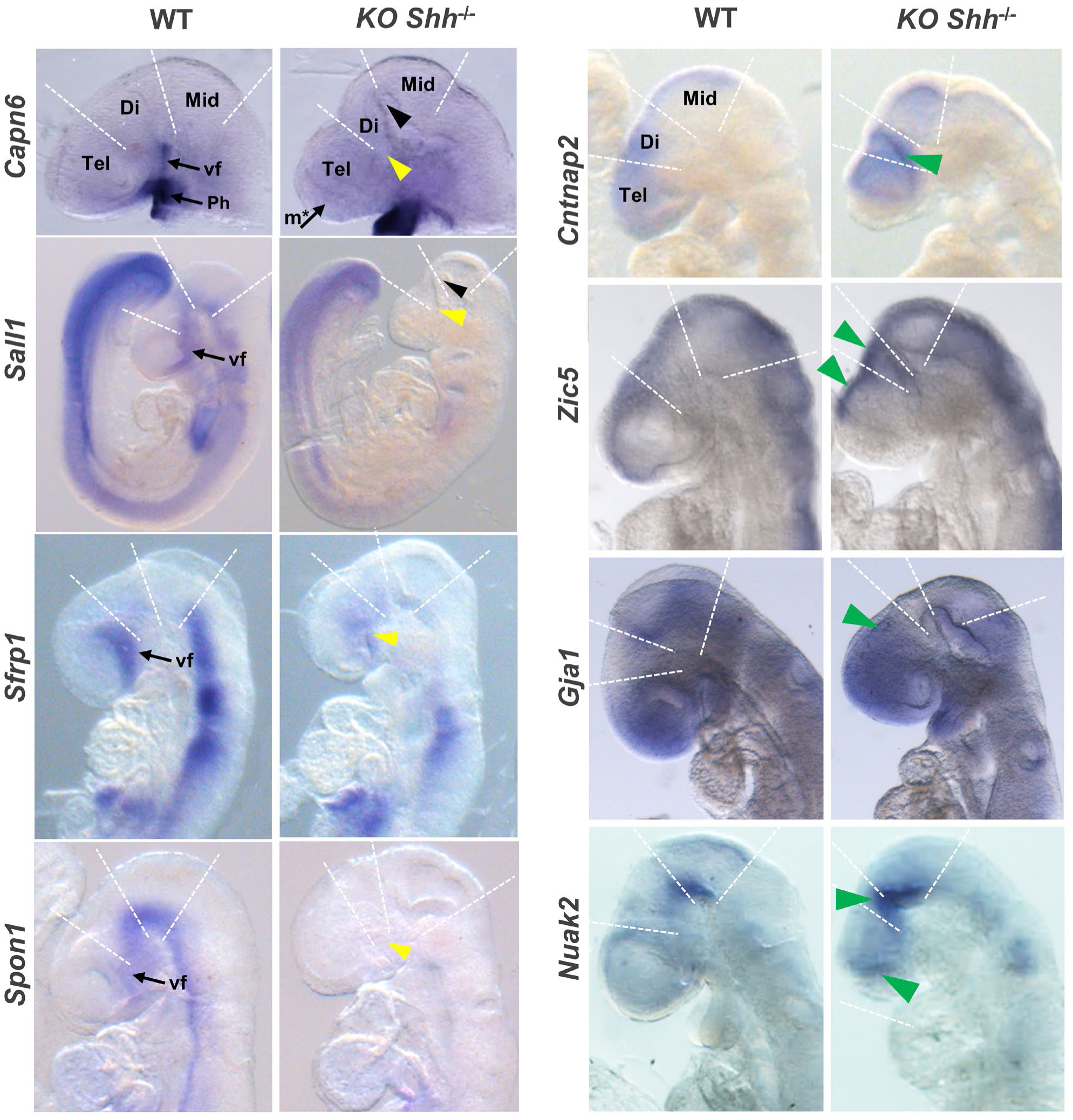
Forebrain expression of selected genes at E9.0 in Wild-type (WT) and Shh knockout (Shh-/-) mouse embryos. Genes analyzed include Capn6, Sall1, Sfrp1, Spon1, Cntnap2, Zic5, Gja1 and Nuak2. Staining was done 3 times for each condition. Black arrowheads indicate the marked ventral constriction at the floor plate level of the diencephalon and midbrain in Shh-/- embryos. Yellow arrowheads highlight region of mRNA expression loss, while green arrowheads indicate gene expression upregulation in mutants. Dashed lines denote boundaries between telencephalon and diencephalon, diencephalon and midbrain, and midbrain and anterior hindbrain.

While *Capn6* has previously been reported to be expressed in the first pharyngeal arch (Tonami et al., 2023), we now demonstrate for the first time its expression in the ventral forebrain as early as E8.25, specifically within the floorplate of the developing forebrain (Supplementary Figure 6). This finding is aligned with its early expression under vAN conditions described in this study (Figure 2). At E9.0. Notably, *Capn6* expression was completely absent in *Shh^-/-^* embryos at specific forebrain areas, including the ventral telencephalon and diencephalon, while it persisted in the pharyngeal arch. Similarly, at E9.0, *Sall1, Sfrp1, Spon1, Rgma, Qki, Pdzrn3, Frzb, Aff2 and Slit1* are broadly expressed throughout the forebrain of WT embryos, but this expression is specifically lost in the anterior ventral region of the forebrain in *Shh^-/-^* embryos (Figure 5 and Supplementary Figure S6). Overall, these findings underscore the essential role of Shh in regulating the expression of these new identified genes during ventral forebrain development in the mouse embryo.

Inversely, we also characterized the expression of *Cntnap2*, *Gja1, Nuak2 and Zic5.* Although these genes are expressed in the developing forebrain at PCW4 in human scRNA (Figure 4), their roles during early vertebrate brain development had not been previously documented. Here, we show that these genes are expressed in the developing dorsal forebrain of mouse embryos, with notably higher expression levels in *Shh^-/-^* embryos.

Whole-mount *in situ* hybridization revealed that *Cntnap2* is tightly regulated during early brain development of WT embryos, with strong expression in the dorsal telencephalon and weaker expression in the dorsal diencephalon and midbrain. Notably, in *Shh^-/-^* embryos at E9.0, *Cntnap2* mRNA levels were increased in the telencephalon and midbrain, with a marked expansion of expression into ventral regions compared to WT embryos (Figure 5, green arrowheads). Similarly, *Zic5*, known for its specific expression along the dorsal neural tube, exhibited significant upregulation in the dorsal forebrain, including both the telencephalon and diencephalon of *Shh^-/-^* embryos. *Gja1*, encoding the gap junction protein connexin 43, was expressed throughout the telencephalon, with its expression extending caudally in *Shh^-/-^* mutants. Interestingly, *Nuak2*, which is involved in microtubule dynamics, is typically localized to the ventral floor plate of the posterior diencephalon and midbrain, showed both increased expression and an anterior expansion of its expression in *Shh^-/-^* embryos (Figure 5, green arrowheads).

These findings provide compelling evidence that SHH signaling orchestrates the precise spatial expression of newly identified genes critical for early forebrain development - some requiring SHH for their ventral expression (e.g., *Capn6, Sall1, Spon1*), while others (*Cntnap2, Zic5, Gja1, Nuak2*) show expanded dorsal expression in its absence. This dual regulation underscores SHH’s essential role in establishing dorsoventral identity and highlights new candidate genes implicated in forebrain patterning.

### Integrative analysis and cross-species validation of novel SHH-associated genes in anterior neuroectoderm development

Considering the promising results obtained from this functional study in the mouse model, we next explored the functional relationships between all the genes in SHH-cluster 1 and SHH-cluster 2 and SHH itself, using the STRING database (Supplementary table 10). Remarkably, the majority of genes identified here - 63% in Shh-Cluster 1 and 94% in SHH-Cluster 2 - had not previously been reported to have a direct (degree 1) topological proximity to SHH.

We created a network representation of the top 100 protein-coding genes from each cluster, selected based on their highest correlation values with SHH within the WGCNA module and their expression in the scRNA-seq dataset at PCW4 (100 out of 545 genes for SHH-cluster 2 and 100 out of 644 for SHH-cluster 1). Additional genes that were tested in mice were also incorporated. Using the STRINGdb interaction database (confidence score ≥ 0.4), we mapped their known interactions with SHH across tissues from various species (Figure 6A). In this network, genes are arranged in concentric circles according to their topological proximity to SHH: genes in the innermost circle (degree 1), directly interact with SHH those in the second interact at degree 2 or 3, and the outermost circle comprises genes with degree 4 interactions or no documented interaction with SHH.

**Figure 6.**
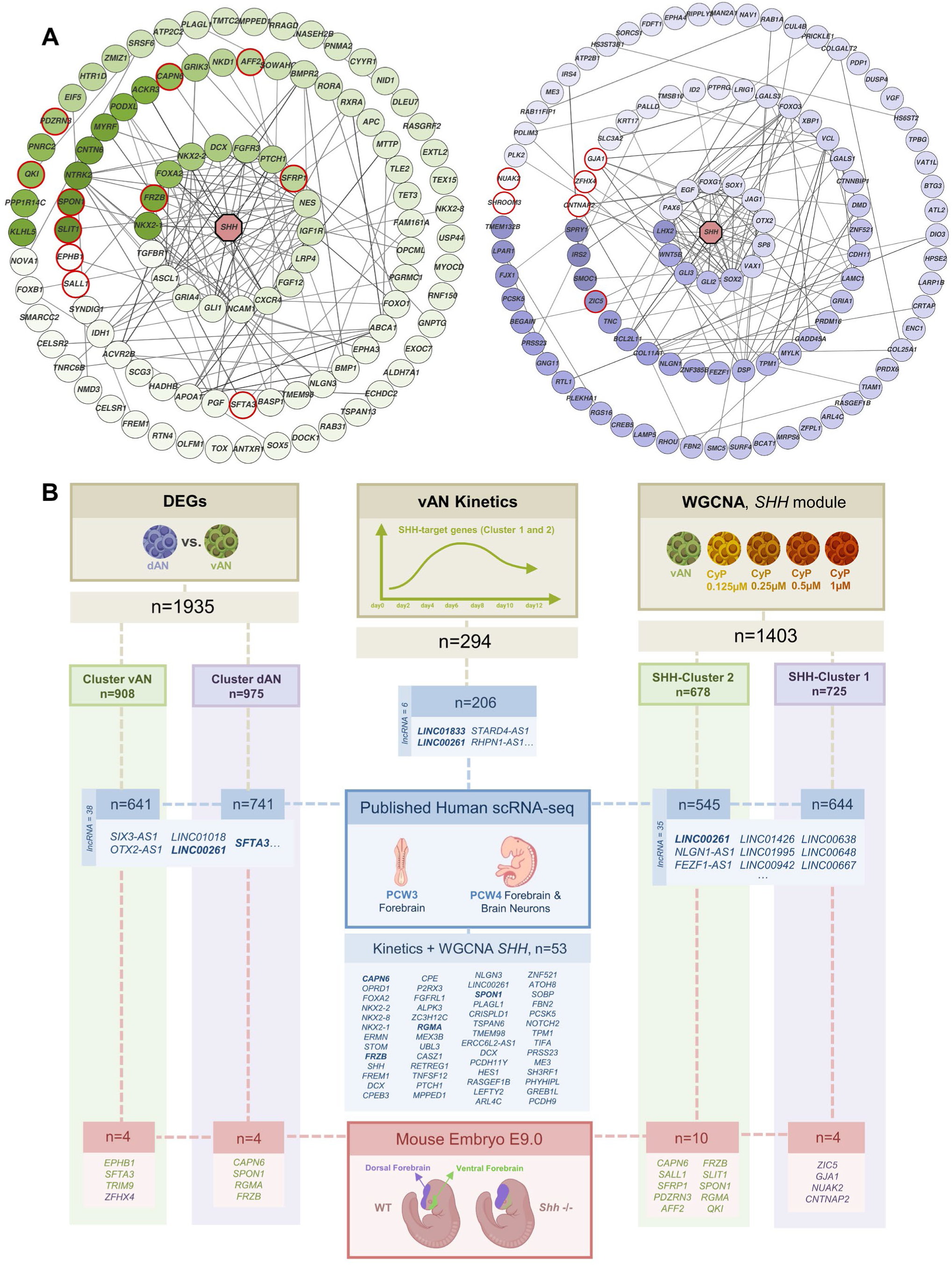
Integrative and cross-species analysis of novel SHH-associated genes during anterior neuroectoderm development. (A) Gene networks based on STRINGdb interactions (confidence score ≥ 0.4), including genes functionally validated in mouse embryos (red nodes) and the top 100 most-co-expressed genes from the WGCNA module containing SHH, expressed in PCW4 human forebrain scRNA-seq data. Nodes represent genes and are color-coded by expression correlation with SHH across LON vAN and CyP samples: green for positive and blue for negative, with color intensity reflecting the strength of correlation. Nodes are arranged in concentric circles according to their topological distance from SHH among these genes: Innermost: 1st-degree; Middle: 2nd-degree to 3rd dege; outermost: 4th degree or higher. Edges thickness indicate STRINGdb interaction confidence score. (B) Schematic overview of the main datasets and analyses integrated in this study. The Brown top boxes indicate the experiment natures: DEGs, subdivided in dAN vs. vAN) from Figure 1; kinetics analysis from Figure 2; WGCNA analysis Figure 4. Each colored box below represents a specific gene set, with the number of genes (n) indicated. Blue boxes highlight genes with expression detected in published human scRNA-seq data (PCW3−PCW4 forebrain and brain neurons, Zeng et al.,2023). Red boxes indicate genes functionally validated in the mouse embryo analysis. Dotted lines and arrows illustrate how these datasets intersect, enabling the identification of both SHH-related and novel genes (including lncRNAs) across multiple model systems.

The inner circle highlights well-established SHH-interacting genes in the forebrain, including *FOXA2*, *NKX2-1*, *PTCH1*, and *HEY1*. Notably, it also features *FRZB* and *SFRP1*, which, to our knowledge, are identified here for the first time as being expressed in the forebrain and regulated by SHH in this tissue during mouse embryonic development. In contrast, genes positioned in the outer circles (interaction degrees 2, 3, or none) are progressively less well characterized regarding their association with SHH. Strikingly, several genes such as *AFF2* and *PDZRN3*, which we demonstrated to be early SHH-responsive genes during mouse brain development, had not previously been linked to SHH signaling in any tissue. Additional genes, including *SALL1, SPON1, RGMA, EPHB1, SLIT1, CAPN6, and AFF2*, also appears in the outer layers (degree 2), further supporting they roles as novel, yet uncharacterized components of the SHH-regulated network during forebrain development.

A similar rationale applies to the 4 genes (*CNTNAP2, ZIC5, GJA1* and *NUAK2*) whose expression was inversely correlated with SHH in our WGCNA analysis (SHH- cluster 1). Although, STRINGdb does not indicate any strong prior association between these genes and SHH signalling in any tissue (Figure 5A), our functional studies in the mouse model revealed that all four are regulated by SHH during early forebrain development. This provides the first evidence that *CNTNAP2*, *ZIC5*, *GJA1*, and *NUAK2* are closely linked to SHH activity in this context, suggesting a previously unrecognized regulatory relationship in early neurodevelopment.

Taken together, these findings suggest that all 545 SHH-dependent genes (SHH- Cluster 1) and all 644 inversely correlated genes (SHH-Cluster 2) identified in this study, and expressed in the prosencephalon of the human embryo at PCW4, may potentially interact with SHH during early forebrain development.

Ultimately, to wrap up our results, we performed a comprehensive integrative analysis of the multiple transcriptomic datasets generated from our human stem cell-derived anterior neuroectoderm models. We systematically cross-referenced these gene lists with scRNA-seq data from human embryonic forebrain tissue at PCW3 and PCW4 (Zeng et al., 2023). This comprehensive approach led to the establishment of gene lists extending key RNA signatures (Figure 6), underscoring the biological relevance of our model. Specifically, we analyzed: (i) 1935 differentially expressed genes (DEGs) under dAN (975 genes) and vAN (908 genes) conditions, (ii) 294 genes identified through temporal pattern analysis, (iii) 678 genes up-regulated by SHH signaling, and (iv) 725 genes down-regulated by SHH signaling. Notably, we also identified several previously uncharacterized long intergenic non-coding RNAs.

This integrative strategy led to the identification of numerous novel genes that had not previously been associated with SHH signaling or forebrain development. To validate the relevance of these findings, we performed functional analyses in mouse embryos, selecting representative genes from each cluster. This cross-species analysis confirmed the biological significance of our *in vitro* results and underscored the evolutionary conservation of early forebrain patterning mechanisms, offering new insights into vertebrate neurodevelopment.

## DISCUSSION

The developmental window between post-conception weeks 3 and 5 marks a critical period in human brain formation, during which the forebrain is first specified. Perturbations in Sonic Hedgehog (SHH) signaling within this interval are known to cause severe neurodevelopmental disorders, including holoprosencephaly. Here, by integrating human iPSC-derived models, transcriptomic profiling, and precise modulation of SHH activity, we established a robust and reproducible *in vitro* system that faithfully recapitulates this critical developmental stage. Importantly, this program is conserved between mouse and human during the initial specification of the ventral neuroectoderm (Brooks et al., 2025). Beyond modeling early human forebrain specification, we identified a set of previously uncharacterized molecular regulators, several of which were validated *in vivo*.

This strategy facilitated a detailed dissection of how SHH activity is interpreted by immediate early genes, highlighting the critical role of temporal dynamics in morphogen signaling. Notably, this study provides the first transcriptomic landscape associated with the morphogenetic, dose-dependent, activity of SHH and underscores the crucial role of temporal dynamics in shaping ventral telencephalon patterning and differentiation. This system, which allows monitoring SHH target gene activation, revealed that immediate early genes act as key decoders of SHH signaling, highlighting temporal modulation as a fundamental layer of SHH function — a principle reminiscent of other pathways such as Ras/Erk.

While several prior studies using brain organoids highlighted SHH’s role in interneuron specification (Cederquist et al., 2019; Bagley et al., 2017; Kim et al., 2019a), our model captures earlier specification events critical for subsequent neuronal differentiation. The deliberate use of a simplified 2D system, prioritizing molecular fidelity over structural complexity, enabled high-resolution dissection of SHH pathway dynamics and provided a valuable resource for future studies of gene regulatory programs governing forebrain regionalization.

The addition of recombinant SHH protein during the early phase of vAN differentiation robustly enhanced ventral commitment, as evidenced by rapid induction of canonical SHH targets such as *PTCH1*, *GLI1*, and *SHH* itself, followed by downstream effectors including *NKX2-1*, *FOXA2*, and *FGF19*, all direct targets of SHH signaling in the mouse ventral midline (Ishibashi et al., 2002; Vincentz, et al., 2005). Among newly identified genes, *FREM1* and *CAPN6* emerged as novel SHH-responsive factors involved in forebrain patterning, both showed expression in early human forebrain (PCW4). *FREM1* is a direct SHH target in mouse neural crest cells, indicating a conserved regulatory role (McLaughlin et al., 2023). Although previously linked to primary cilia formation but not brain development (Tonami et al., 2007), CAPN6 is shown here to be closely associated with SHH signaling during vAN differentiation as in the mouse ventral forebrain; notably, its expression was abolished upon SHH pathway inhibition. Identifying these two genes marks a key advance in understanding brain development and suggests that many of the other genes highlighted in this study may also be involved in early forebrain development.

Several other genes associated with neurodevelopment, such as *SFRP1*, *AFF2*, *QKI*, and *PDZRN3 (*Oliver et al., 2003; Baizabal et al., 2018; Zou et al., 2022; Chénard and Richard, 2008), but not previously associated with SHH, were identified, revealing an unanticipated breadth of SHH-regulated networks from the earliest stages of forebrain development.

Interestingly, a subset of genes, including *CNTNAP2*, *GJA1*, and *NUAK2* displayed increased expression as SHH signaling declined, suggesting that SHH initially acts as a repressor of specific neurodevelopmental programs. This repressive role was confirmed in Shh^⁻/⁻^ mouse embryos. These genes have known roles in cortical development and neurodevelopmental disorders, supporting the importance of early SHH-mediated repression in orchestrating cortical fate specification (Saint-Martin et al., 2018; Iacobas et al., 2007; Blazejewski et al., 2021).

In addition to protein-coding genes, our study highlighted the regulation of multiple lncRNAs by SHH signaling. Notably, *SFTA3*, *LINC01833*, and *LINC00261*, which exhibit spatial and functional associations with key neurodevelopmental regulators (Herriges et al., 2017; Swarr et al., 2019). These findings underscore the emerging significance of lncRNAs as regulatory elements in vertebrate forebrain development.

From a translational perspective, beyond the characterization of all these genes as potential novel biomarkers of neurodevelopmental pathologies, our model offers a valuable platform for studying human neurodevelopmental disorders. Leveraging hiPSC technology, we established a system capable of generating ventral anterior neuroectoderm cells amenable to transcriptomic analyses.

This system facilitated the identification of transcriptomic signatures correlating with graded SHH pathway inhibition, representing the first such report for Holoprosencephaly-related defects. Recent studies suggest that transcriptomic signatures of complex disorders offer a promising avenue for improving the interpretation of genotypes (Curry et al., 2021; Yépez et al., 2022). Likewise, patient-derived iPSCs, combined with our robust differentiation model and reference signatures, will provide an opportunity to stratify holoprosencephaly cases molecularly and functionally validate disease-associated alleles. Given the correlation between SHH pathway disruption and holoprosencephaly severity observed in animal models (Hong and Krauss, 2018; Mercier et al., 2013), these transcriptomic signatures may also serve as prognostic indicators. Our data thus establish the biological relevance of this model for studying SHH-dependent forebrain defects, with implications extending to other Hedgehog-related disorders such as ciliopathies (Andreu-Cervera et al., 2019).

In summary, this work significantly advances our understanding of the temporal regulation of SHH signaling during early human forebrain development, identifies novel molecular players and biomarkers, and establishes a reference framework for improving the diagnosis of SHH-related neurodevelopmental disorders. The iPSC-derived models described here sets a new benchmark for disease modeling and personalized medicine approaches.

## Lead contact

Valérie DUPÉ –

## Materials availability

This study did not generate new unique reagents.

## Resource availability

Sequencing data are currently in the process of being submitted to https://ega-archive.org and can be provided upon request. All original code has been deposited on GitHub and is publicly available at https://github.com/GeDiNe-Lab/panasenkava_2024 as of the date of publication. Any additional information required to reanalyze the data reported in this paper is available from the lead contact upon request

## Supporting information

Supplementary Tables Part 1

Supplementary Tables Part 2

Supplementary Tables Part 3

Supplementary Tables Part 4

## Acknowledgments

The authors acknowledge the network FHU GenoMeds and the BIOSIT facilities (Univ Rennes, CNRS, Inserm, Biosit UAR 3480 US_S 018, Rennes, France). This work was funded by the Agence Nationale de la Recherche (ANR-22-CE17-0064 and ANR-24- CE16-2483-01).

## Author contributions

Conceptualization, V.P and V.D; Methodology, V.P, H.G, F.D, M.D.T and V.D; Software J.G, W.C, A.M and M.D.T; Validation V.P, H.G, Y.V, E.J and L.D; Formal analysis V.P, J.G, A.M, Y.V and L.D; Investigation V.P and J.G; Data Curation J.G, W.C and M.D.T; Writing - Original Draft, V.P and V.D; Writing - Review & Editing, V.P, V.R, A.C and V.D; Visualization, V.P and J.G; Supervision V.D; Funding acquisition V.D.

## Supplementary Figures

**Figure S1:** Validation of control hiPSCs lines (LON71 and WTC11) used in the differentiation study

**Figure S2:** Temporal dynamics of dorsal and ventral anterior neuroectoderm differentiation from human iPSCs (LON cell line).

**Figure S3:** Assessment of cell identity at the mid-point of the differentiation protocol (day 6) in LON line.

**Figure S4:** Supplementary analysis of SHH pathway inhibition through Cyclopamine gradient in anterior neuroectoderm differentiation.

**Figure S5:** Gene’s selection expression verification in human brain scRNA-seq data.

**Figure S6:** Expression of several newly characterized genes in the forebrain at E9.0 and E8.25 of Wild-type (WT) and E9.0 of *Shh* mutants (*Shh* ^-/-^).

## Supplementary Tables titles

**Supplementary table 1 (Figure 1E):**

Differential expression analysis between vAN and dAN samples at day 12 (LON71 lineage)

**Supplementary table 2A and 2B (**Figure 1F):

GO terms DEGs vAN/dAN day 12

**Supplementary table 3A and 3B (**Figure 2C):

PCA associated genes (heatmap) for WTC and LON

**Supplementary table 4A-4H (Figure S2B):**

GO terms kinetic (WTC and LON)

**Supplementary table 5 (**Figure 2D):

DEGs at day 2 between vAN and dAN

**Supplementary table 6A-6E (**Figure 2):

Differential expression analysis between time points for vAN

**Supplementary 6F-6J (**Figure 2):

Differential expression analysis between time points for dAN

**Supplementary table 7A and 7B (**Figure 2H **& S2):**

degPattern results

**Supplementary table 8A-8E (**Figure 2H):

degPattern focus clusters with SHH STRINGdb networks

**Supplementary table 9A and 9B (**Figure 4 **& S4):**

SHH WGCNA module for LON and WTC

**Supplementary table 10 (**Figure 6):

SHH WGCNA module (LON) STRINGdb network

**Supplementary table 11 (**Figure 6):

Figure genes recapitulating table

## METHODS

### Maintenance of human iPSCs

2 hiPSC lines were used in this study: LON71.019 (<u>Gaignerie *et al*., 2018</u>; generated in the iPSC core facility of the University of Nantes), WTC11 (UCSFi001-A, commercial line from Conklin Lab, Gladstone/UCSF). All studies involving hiPSC were conducted in accordance with approved protocols. All lines were tested for normal karyotype, negative mycoplasma, appropriate stem cell/colony morphology and genetic integrity (Infinium Core-24 Kit, Illumina). For culture maintenance cells were grown at 37°C and expanded in monolayer on Matrigel coated (CORNING, #354277, 1:100 ratio with DMEM/F12) plastic cell cultures multiwell plates in mTeSR™ Plus medium (STEMCELL, #100-0276) with daily medium change, until reaching an approximate confluence of 80% before passage. For passage, cells were dissociated for replating using StemMACS™ Passaging Solution XF (MILTENYI BIOTEC, #130-104-688). Y- 27632 (STEMCELL, #72304, 1:1000 ratio) was added in cell medium after each passage and thawing to preserve cells from dissociation-induced apoptosis and increase the survival.

### Differentiation of hiPSCs in Anterior Neuroectoderm

Before inducing neural differentiation, cells were detached with Accutase (STEMCELL, #07920) to make a single-cell suspension, then replated on Matrigel coted plates (1:50 ratio with DMEM/F12) and grown until approximately 95%-100% confluence, covering most of the surface area of the culture plate wells. 12 days differentiation protocol is based on <u>Tchieu et al., 2017</u> (see detailed protocol for “Neuroectoderm differentiation”). In brief, 2 culture media were used: KSR (DMEM KnockOut™ #10829018 Thermo Fisher Scientific, Knockout Serum Replacement #10828028 Thermo Fisher Scientific, Penicillin-Streptomycin #15140122 Gibco™, L-Glutamine 200 mM #25030081 Thermo Fisher Scientific, MEM Non-Essential Amino Acids 100x #11140050 Thermo Fisher Scientific, 2-mercaptoethanol 55 mM #21985023 Thermo Fisher Scientific) and N2 (DMEM/F12 #11320033 Thermo Fisher Scientific, 2- mercaptoethanol 55 mM #21985023 Thermo Fisher Scientific, Sodium Bicarbonate #S5761-500G Sigma-Aldrich, D-(+)- Glucose #G8270 Sigma-Aldrich, Progesterone #P8783-1G Sigma-Aldrich, N2 Supplement-B #07156 STEMCELL). 3 chemicals were added to induce neural anterior fate: LDN193189 #72147 500nM, SB431542 #72232 10µM, XAV939 #72672 5µM, STEMCELL. To induce ventralisation and generate ventral anterior neuroectoderm 500ng/ml of human recombinant SHH (C24II, #78065, STEMCELL) was added to media mix from day 0 to day 6. For gradual SHH pathway inhibition 1µM, 0.5µM, 0.25µM or 0.125µM of CyP (#C4116, Sigma-Aldrich) was added to media mix after ventralisation from the day 6 to the day 12. After differentiation ends cells were detached with Accutase, pelleted and stored at-80°C prior ARN extraction.

### RNA extraction

Total RNA was extracted from cell pellets using the Maxwell® 16 miRNA Tissue Kit (#AS1460, Promega) on the AS2000 Maxwell® 16 Instrument (IRSET, Rennes, France). The concentration of RNA was determined using spectrophotometry method (Nanodrop Microvolume Spectrophotometer). Subsequently, supplementary DNase treatment was performed on the samples, using 1 unit of DNase I (#AMPD1, Sigma-Aldrich) per microgram of RNA before RT-qPCR assay.

cDNA was synthesized using High-Capacity cDNA Reverse Transcription Kit (#4368814, Applied Biosystems™) according to the manufacturer’s protocol in a total reaction volume of 40-60 μl. Quantitative real-time PCR was performed using PowerUp™ SYBR™ Green Master Mix (#A25742, Applied Biosystems) according to the manufacturer’s protocol in QuantStudio™ 7 Flex Real-Time PCR System. The cycling parameters were as follows: 2min at 50 °C, 10 min at 95 °C for initial denaturation, followed by 40 cycles of 15 s at 95 °C, 1min at 60 °C, and 15s at 95 °C. Gene specific primer sequences are listed in Table 1. Finally, data were analyzed using ΔΔCt-quantification method. All analyses were performed with three technical replicates per plate.

**Table 1.**
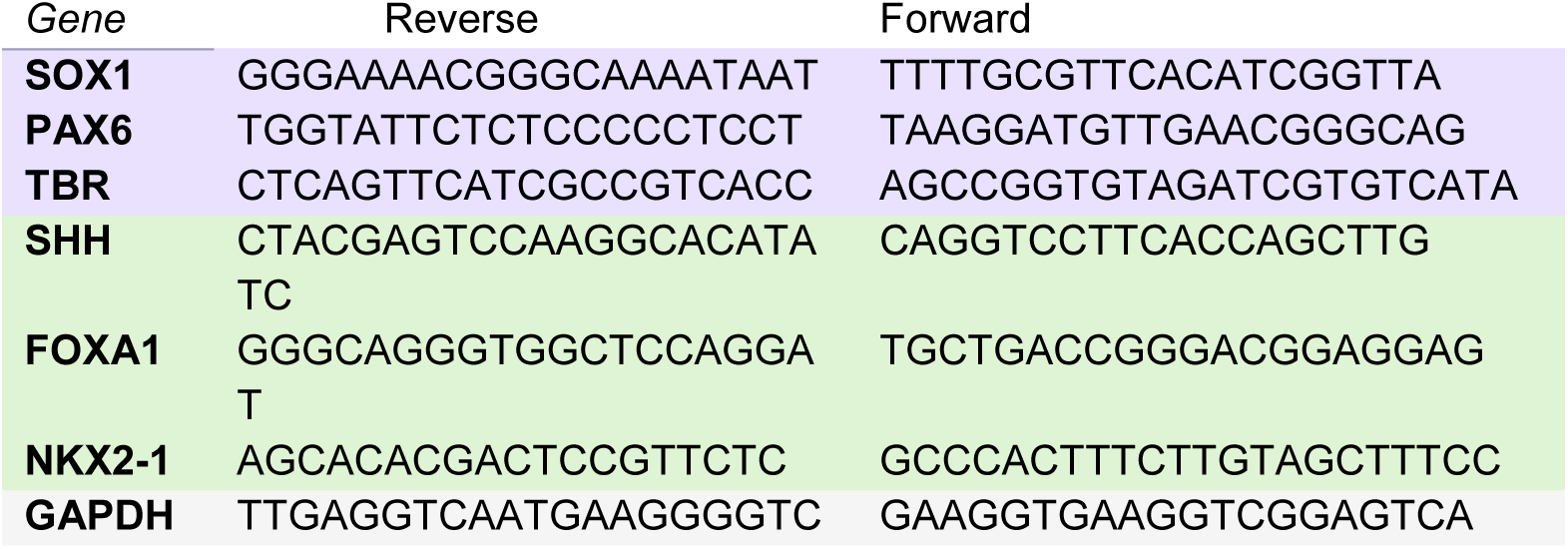
List of primers used for RT-qPCR assay.

### RT-qPCR gene expression study

#### Immunofluorescence Assays

The cells grown on the Matrigel coated coverslips or imaging chambers-slides (Lab-Tek™ II Chamber Slide #154534PK Thermofisher), then were fixed in 4% paraformaldehyde for 15 minutes at room temperature. Triton X-100 (0.20%) was used for permeabilization, followed by incubation with blocking reagent BSA 3% (Bovine Serum Albumin Fraction V, Roche) in PBS1X for 1h. Primary antibodies were diluted in a BSA 1% PBS1X and incubated overnight at 4°C in a humidified chamber. After three PBS washes (5 min each), secondary antibodies were added for 2h at room temperature. Cells were washed 3 times with PBS and then DAPI was used to counterstain nuclei (5748, 1:5000, R&Dsystems). List of all antibodies used in experiment is available in Table 2. Cells or slices were then mounted in ProLong™ Gold Antifade Mountant (#P36934, Invitrogen) or VECTASHIELD® Hardset Antifade Mounting Medium with DAPI (#H-1500-10, Vector Laboratories), then imaged by a confocal microscope (Leica TCS SP8, Biosit, Rennes, France).

**Table 2.** List of Antibodies used for Immunofluorescence assay.

| **Antibody** | **Reference** | **Supplier** |
| --- | --- | --- |
| Human TBR2/EOMES | MAB6166-SP | R&Dsystems |
| Human HNF-3 beta /FoxA2 | AF2400-SP | R&Dsystems |
| Human TTF-1/NKX2-1 | MAB94581-SP | R&Dsystems |
| Human SOX1 | AF3369-SP | R&Dsystems |
| Human/mouse SHH (N-Ter) | AF464-SP | R&Dsystems |
| Human/mouse EMX2 | AF6470 | R&Dsystems |
| Donkey Anti-Sheep IgG NorthernLights™ NL637 | NL011 | R&Dsystems |
| Donkey Anti-Goat IgG NorthernLights™ NL637 | NL002 | R&Dsystems |
| Donkey Anti-Rabbit IgG NorthernLights™ NL637 | NL005 | R&Dsystems |
| DAPI | 5748 | R&Dsystems |
| Anti-Mouse 637 | NL008 | R&Dsystems |
| Anti-Goat 493 | NL003 | R&Dsystems |
| Anti-Sheep 557 | NL010 | R&Dsystems |
| Alexa Fluor™ 647 | A21247 | Invitrogen |
| Alexa Fluor™ 488 | A11034 | Invitrogen |
| Alexa Fluor™ 546 | A11035 | Invitrogen |
| Human PAX6 | sc-81649 | SANTA CRUZ |
| ANTI-MOUSE m-IgG2b BP-CFL 488 | sc-542745 | SANTA CRUZ |
| ANTI-MOUSE m-IgGk BP-CFL 488 | sc-516176 | SANTA CRUZ |

#### Mice

The Shhtm1Amc/J mouse strain was purchased from the Jackson Laboratory (Bar Harbor, ME, USA) and maintained on a C57BL/6J background (Janvier Labs, France). Embryos stages used for this study were between E8.5 and E9 day post coitum. To score embryonic stages more precisely, somite numbers were counted.

#### In situ hybridization

All embryos were fixed in 4% PFA/PBS at 4°C overnight, rinsed and processed for whole-mount ribonucleic acid (RNA) in situ hybridization. Probes was created by PCR amplification, using specific primers and cDNA bank of mice testis. Then DNA obtain was cloned on pCRII-TOPO TA vector (452640, Invitrogen) and Digoxigenin probes were synthesized using Digoxigenin RNA labeling kit (Roche Cat. No. 11 175 025 910). Whole mount *in situ* hybridization of embryos was performed as previously described (Ratié *et al*. 2013)

#### Ethics statement

All experiments with mice were carried out in accordance with the European Communities Council Directive of November 24, 1986. VD, as the principal investigator in this study, was granted the accreditation 35–123 to perform the reported experiments and the experimental procedures were authorized by the French Ministry of Research committee C2EA-07 under the protocol APAFIS#3532- 2016010514029297.

#### RNA-sequencing & Data analysis

The RNA was sent to Integragen® sequencing platform for libraries preparation and sequencing according to 3’RNA-Seq sequencing Protocol (NovaSeq, 2 * 100 bases). FastQ files were recovered and analyzed with a standardized pipeline available at (github link).

#### RNA-sequencing data processing

RNA sequencing pipeline starts with a trimming step adapted to 3’RNA-Seq protocol using Cutadapt. A first trimming of the reads 1 (R1) allows to keep only the 26 first nucleotides in 5’. In parallel, a specific trimming of read 2 (R2) is performed with: trimming of the 4th first base in 3’, filtering on quality (minimum quality phred score = 20), removing of polyA tails and keeping of reads with a minimum length of 50 nucleotides. The R1 trimming keeps only the information about the UMI and the R2 trimming keeps only the sequence of interest. The first alignment of R2 reads is performed using STAR. Then, the output bam files are sorted by read’s name with sambamba and the UMIs of the R1 are assigned to the R2 reads in the bam file with fgbio (AnnotateBamWithUmi). The mate-pair information is verified by Picard (FixMateInformation) and the output bam files are sorted by coordinates. The duplicates UMIs are removed with UmiAwareMarkDuplicatesWithMateCigar from Picard. The annotated bam file is sorted with the UMI and CIGAR codes using the coordinates with sambamba. Then, recovery of annotated fastq files is performed by bamtofastq from bedtools. Finally, the quantification is performed by an alignment of the annotated fastq on the human genome reference (version hg38) with STAR (quantMode GeneCounts).

#### Counts normalization

Counts were normalized by variance stabilizing transformation using the vst function from the DESeq2 package on DESeqDataSet object. Normalization was done by considering DESeqDataSet objects design with different covariates depending on the analyzed data. Covariates were the following: type (dorsal, ventral or CyP), hiPSCs lineage, differentiation day, CyP dosage.

#### Principal component analysis (PCA)

PCA analyses were performed on normalized data by variance stabilizing transformation without covariates on the top 3000 genes with highest variance across samples. We used prcomp function from the stats package. Association between PCs and experimental variables were tested through a one-way ANOVA using the aov function from the stats package. Gene selected for Figure 2C were selected as genes in *(A ∪ B) \ C* where *A*, *B*, and *C* are the top 1000 most associated genes with PC1, PC2, and PC3 respectively.

#### Differential gene expression analysis

Differential gene expression analyses were performed using the DESeq function from the DESeq2 packages with default parameters. Significant differentially expressed genes were defined as those with FDR < 0.01 (Benjamini-Hochberg multiple testing correction) and |log2FC| ≥ 1. We also defined genes with FDR < 0.01 and |log2FC| ≥ 2 as highly differentially expressed genes.

#### Expression patterns

Genes expression patterns were computed using the degPatterns function from the DEGReport package with reduce parameter set to True, nClusters = 10 and other parameters as default. degPatterns compute pair-wise gene expression using cor.test R function with kendall method then turn this correlation matrix to distance matrix. gene clustering is performed using cluster::diana(). Tree cutting is done using clustering divisive coefficient.

#### Hierarchical culstering

Hierarchical clustering were performed using the fastcluster::hclust() function with complete method on a dissimilarity matrix from the selected genes and samples.

#### WGCNA analysis

We perform WGCNA on the normalized data using the top 50% genes with highest statistical variance (10165 and 9151 genes for LON71 and WTC lineage respectively). Adjacency matrix was computed as an unsigned network with a power of 12 for LON71 lineage and 13 for WTC lineage. Topological overlap matrix was computed using TOMsimilarity with default parameters and genes were clustered using hierarchical clustering. dynamic tree cut was performed with the following parameters: deepSlit = 2, pamRespectsDendro = False, minClusterSize = 30 (we kept other parameters as default). Modules with an eigengenes correlation higher than 0.75 were merged using the mergeCloseModules function.

#### Gene Ontology enrichment analyses

Gene Ontology (GO) enrichment analyses were done using the enrichGO function from the clusterProfiler package with default parameters. We kept Biological process term with a p-value < 0.05.

#### scRNA-seq-data analysis

We used scRNAseq data from Zeng et al., 2023. We used cells from central nervous system at PCW3, PCW4 and PCW5 according to the author’s annotation. UMAP were computed with the Seurat R package. We used RunUMAP function using dimensions 1 to 10 from the results of the RunPCA function with 2000 variable features.

#### Statistics

Statistical analyses were conducted using GraphPad Prism 8.0.1 for the Windows OS. Additional information and the number of samples (n) for each experiment can be found in the respective figure legends. The analysis of the data employed a one-way ANOVA statistical test. The data is presented as the mean ± standard error of the mean (SEM), and a P-value less than 0.05 was considered statistically significant. For the rest of the results, significance levels are indicated as follows: *p < 0.05, **p < 0.01, ***p < 0.001.

