## Supplementary Information for "Mapping the temporal and functional landscape of Sonic Hedgehog signaling reveals new insights into early human forebrain development"

### Supplementary Figures List:

**Figure S1:** Validation of control hiPSCs lines (LON71 and WTC11) used in the differentiation study

**Figure S2:** Temporal dynamics of dorsal and ventral anterior neuroectoderm differentiation from human iPSCs (LON cell line).

**Figure S3:** Assessment of cell identity at the mid-point of the differentiation protocol (day 6) in LON line.

**Figure S4:** Supplementary analysis of SHH pathway inhibition through Cyclopamine gradient in anterior neuroectoderm differentiation.

**Figure S5:** Gene's selection expression verification in human brain scRNA-seq data.

**Figure S6:** Expression of several newly characterized genes in the forebrain at E9.0 and E8.25 of Wild-type (WT) and E9.0 of *Shh* mutants (*Shh* $^{-/-}$).

### Supplementary Tables titles List:

**Supplementary table 1 (Figure 1E):**
Differential expression analysis between vAN and dAN samples at day 12 (LON71 lineage)

**Supplementary table 2A and 2B (Figure 1F):**
GO terms DEGs vAN/dAN day 12

**Supplementary table 3A and 3B (Figure 2C):**
PCA associated genes (heatmap) for WTC and LON

**Supplementary table 4A-4H (Figure S2B):**
GO terms kinetic (WTC and LON)

**Supplementary table 5 (Figure 2D):**
DEGs at day 2 between vAN and dAN

**Supplementary table 6A-6E (Figure 2):**
Differential expression analysis between time points for vAN

**Supplementary 6F-6J (Figure 2):**
Differential expression analysis between time points for dAN

**Supplementary table 7A and 7B (Figure 2H & S2):**
degPattern results

**Supplementary table 8A-8E (Figure 2H):**
degPattern focus clusters with SHH STRINGdb networks

**Supplementary table 9A and 9B (Figure 4 & S4):**
SHH WGCNA module for LON and WTC

**Supplementary table 10 (Figure 6):**
SHH WGCNA module (LON) STRINGdb network

**Supplementary table 11 (Figure 6):**
Figure genes recapitulating table

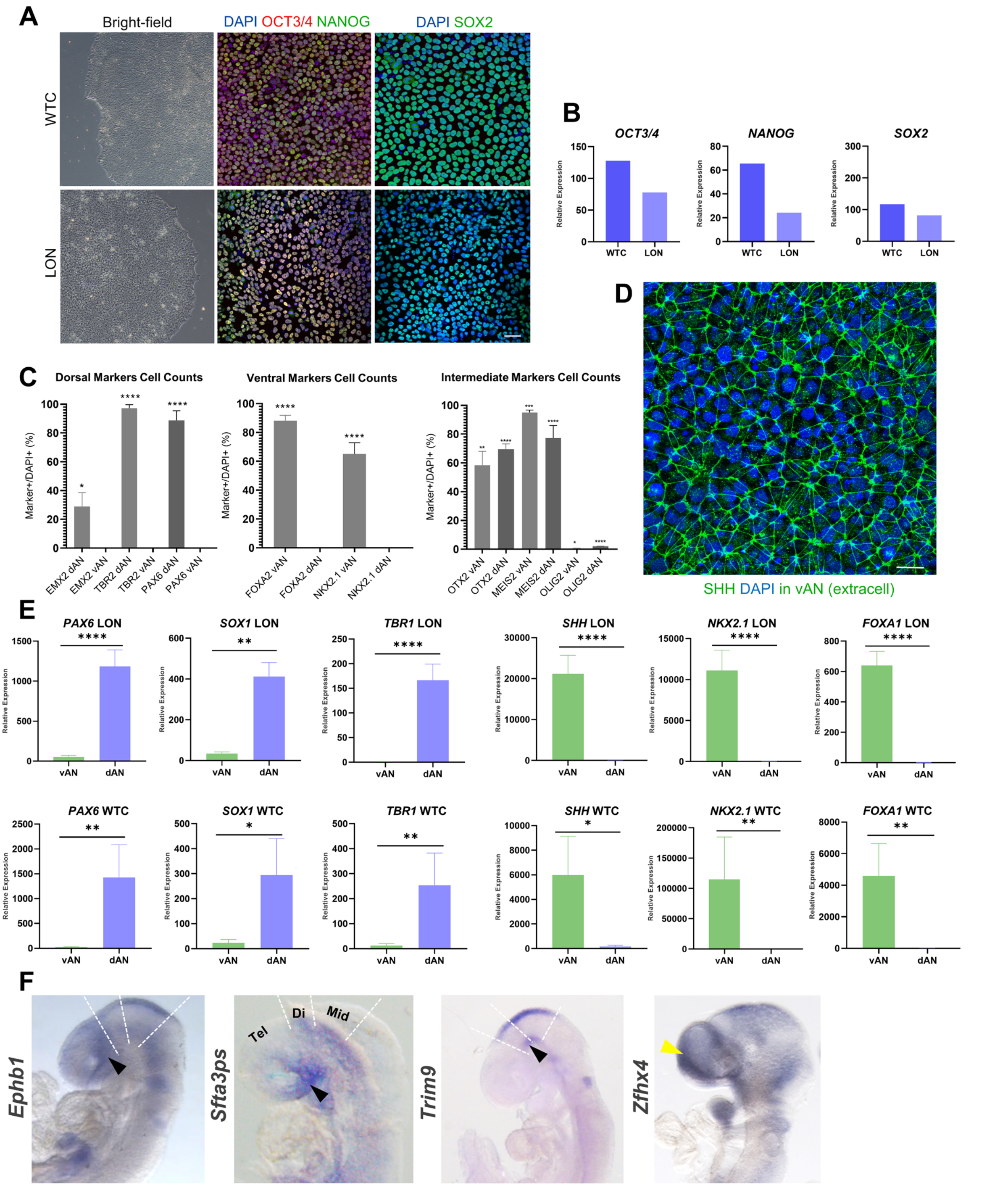


**Figure S1: Validation of control hiPSC lines (LON71 and WTC11) used in the differentiation study.**

**(A)** Representative images of the LON and WTC hiPSC lines. Bright-field images (top row) illustrate typical colony morphology, showing uniform and undifferentiated cellular morphology consistent with pluripotency. Immunofluorescence staining (middle and bottom rows) confirms expression of pluripotency markers OCT3/4, NANOG, and SOX2 in both lines, with DAPI staining marking cell nuclei (scale bar: 40 µm).

**(B)** Quantitative RT-PCR analysis of pluripotency marker expression (*OCT3/4, NANOG*, and *SOX2*) in LON and WTC hiPSC lines. Expression levels are represented as relative expression normalized to endogenous controls, confirming robust pluripotency marker expression in both cell lines, validating their suitability for directed differentiation.

**(C)** Quantification of positive cells for dorsal, ventral, and intermediate markers in both dAN and vAN conditions at day 12 in LON line, shown as a percentage of total DAPI-positive cells. Dorsal markers (e.g., PAX6, TBR1) exhibit higher positivity in dAN, whereas ventral markers (e.g., FOXA2, NKX2.1) show higher positivity in vAN. Intermediate markers, representing shared or transitional phenotypes, also vary according to differentiation condition. Values are presented as mean ± SEM, with significance levels marked as *$p < 0.05$, **$p < 0.01$, ***$p < 0.001$, ****$p < 0.0001$ (t-tests), highlighting the distinct regional identities achieved in each condition.

**(D)** Immunofluorescent staining for extracellular Sonic Hedgehog (SHH) in ventral anterior neuroectoderm (vAN) cultures at day 12 of differentiation of LON line. SHH protein localization is visualized in green, and nuclear DAPI staining in blue highlights cell nuclei, illustrating extracellular SHH presence in the vAN culture, crucial for establishing ventral identity.

**(E)** Relative expression levels of dorsal (*PAX6, SOX1,* and *TBR1*) and ventral (*SHH*, *NKX2.1,* and *FOXA1*) marker genes measured by RT-qPCR at day 12 of differentiation in two hiPSCs lines, LON and WTC, in dorsal (dAN) and ventral (vAN) conditions. Each gene's expression level is normalized and compared between vAN and dAN, with 5 replicates for LON and 2 replicates for WTC. The results confirm a significant upregulation of dorsal markers in dAN and ventral markers in vAN. Statistical significance is indicated as follows: *$p < 0.05$, **$p < 0.01$, ***$p < 0.001$, ****$p < 0.0001$, based on t-tests.

**(F)** Expression of several newly characterized genes in the forebrain at E9.0 of Wild-type (WT) . The genes analyzed include *Ephb1, Sfta3ps (lincRNA), Trim9, Zfhx4*. Black arrowheads indicate the pronounced ventral constriction at the level of the floor plate of the diencephalon and midbrain of *Shh* -/- embryos. Yellow arrowheads indicate the loss of mRNA expression, while green arrowheads indicate the upregulation of gene expression in *Shh* -/- embryos. Dashed lines represent the boundaries between telencephalon and diencephalon, the diencephalon and midbrain, and the midbrain and anterior hindbrain.

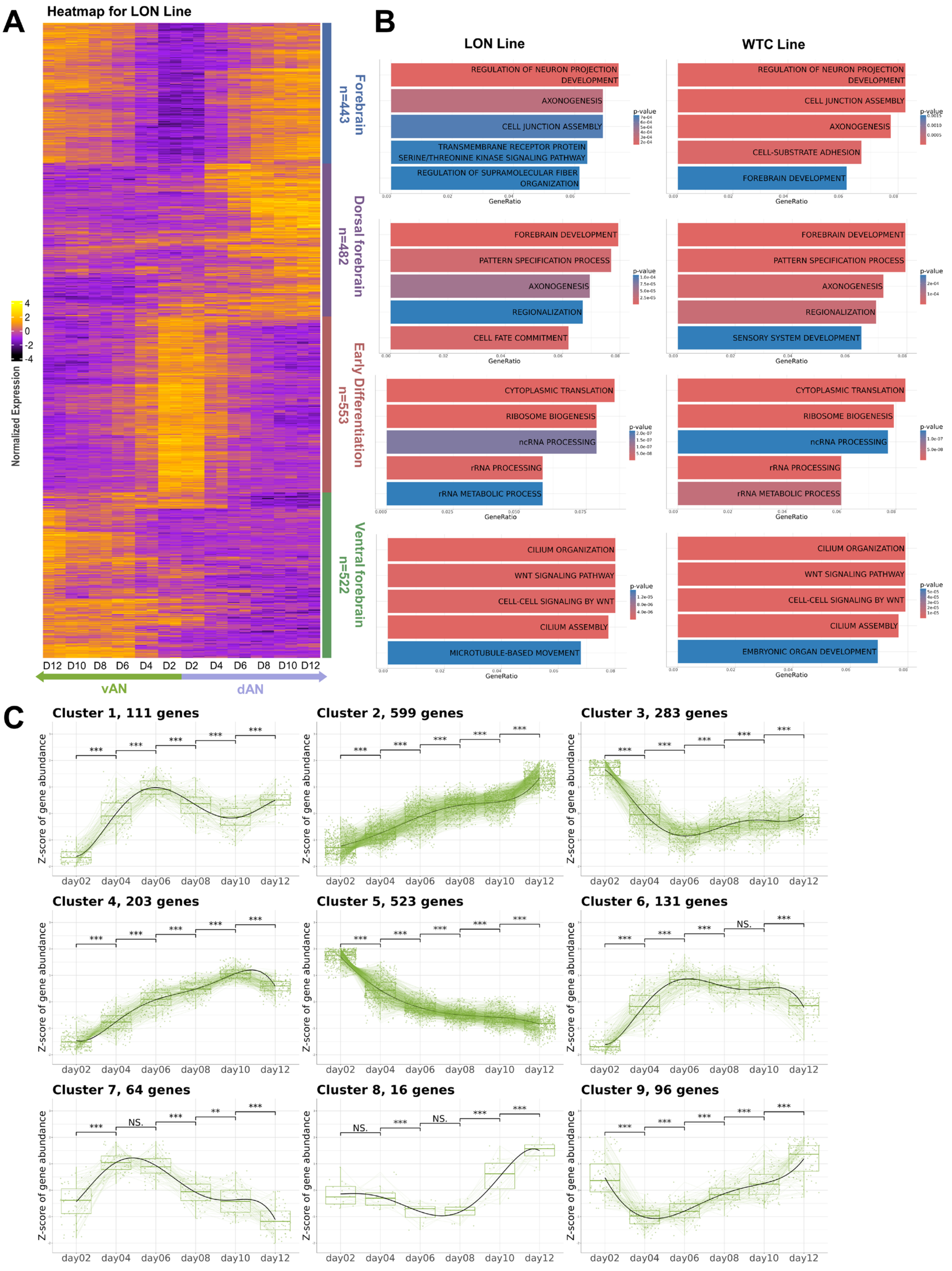


**D**

| STRING DataBase | Cluster 1 (n=111) | Cluster 2 (n=599) | Cluster 6 (n=131) |
|---|---|---|---|
| *SHH*-related | *DCX, DKK1, FEZF2, FGF9, FRZB, LEF1, NOTCH2 (n=7, 6.3%)* | *NKX2-1, BMP2, BMP4, BMP7, DKK3, EFNB2, EGFR, EYA1, FABP7, FN1, FOXA1, LRP2, NTN1, PITX2, POU3F2, TBX1…(n=35, 5.8%)* | *SIX6, SOX3, CHRD, FGF10, FOXA2, GPC3, HEY1, HHAT, PTCH1, RNF3 (n=10, 7.6%)* |
| Novel genes | ***TSPAN6, TMEM98, CAPN6, SLC4A11, ZFHX4, EGR1, MXD4, SOX4, GRIA3, ZNF436, AMOT, PALLD, NKX2-8… (n=100, 90%)*** | ***SPON1, SLIT2, NLGN3, CTNNA3 LHFPL6, PLCE1, DDC, TRIM9, PLCL1, FOXP1 … (n=556, 92.8%)*** | ***SHOC1, SALL2, CHST14, NLGN1, SDR9C7, TMC7, PDE7B, HOPX, RAB37, CLCF1…(n=117, 89%)*** |
| Novel lncRNA | ***LINC02981, LINC00261...(n=4, 3.6%)*** | ***LINC02487, LINC01252… (n=8, 1.3%)*** | ***LINC00954, LINC02984…(n=4, 3%)*** |

**Figure S2: Temporal dynamics of dorsal and ventral anterior neuroectoderm differentiation from human iPSCs (LON cell line).**

**(A)** Heatmap showing scaled normalized expression of the top 1000 most associated genes with the two first principal component of the PCA respectively (Pearson correlation between principal component eigenvector and gene expresion across samples) for samples from the LON cell lines. Hierarchical clustering was applied to genes, while sample where mannually organized based on type and timepoints. Clusters were defined using the cutree R function with parameter k = 4. Gene clusters are annotated based on anterior forebrain markers (n=467), dorsal forebrain markers (n=526), early differentiation markers (n=547), and ventral forebrain markers (n=460).

**(B)** Top 5 most enriched Gene Ontology biological process terms in each gene cluster. Panels are associated to anterior forebrain, dorsal forebrain, early differentiation and ventral forebrain clusters respectively from top to bottom (First column for LON line, second column for WTC line).
**(C)** Clusters from DEGpattern results for genes differentially expressed between at least 2 consecutive timepoints on LON samples. Each plot shows the z-score–normalized expression trajectories over time for the genes within each cluster (green lines), with the average trend depicted in bold. Statistical significance of differences among time points is indicated by asterisks (***$p < 0.001$).

**(D)** Genes from Cluster 1, Cluster 2, and Cluster 6 (from Figure S2C) were analyzed using the STRING database to identify known or predicted relationships to the Sonic Hedgehog (SHH) pathway ("SHH-related"), as well as to highlight novel genes and lncRNAs.

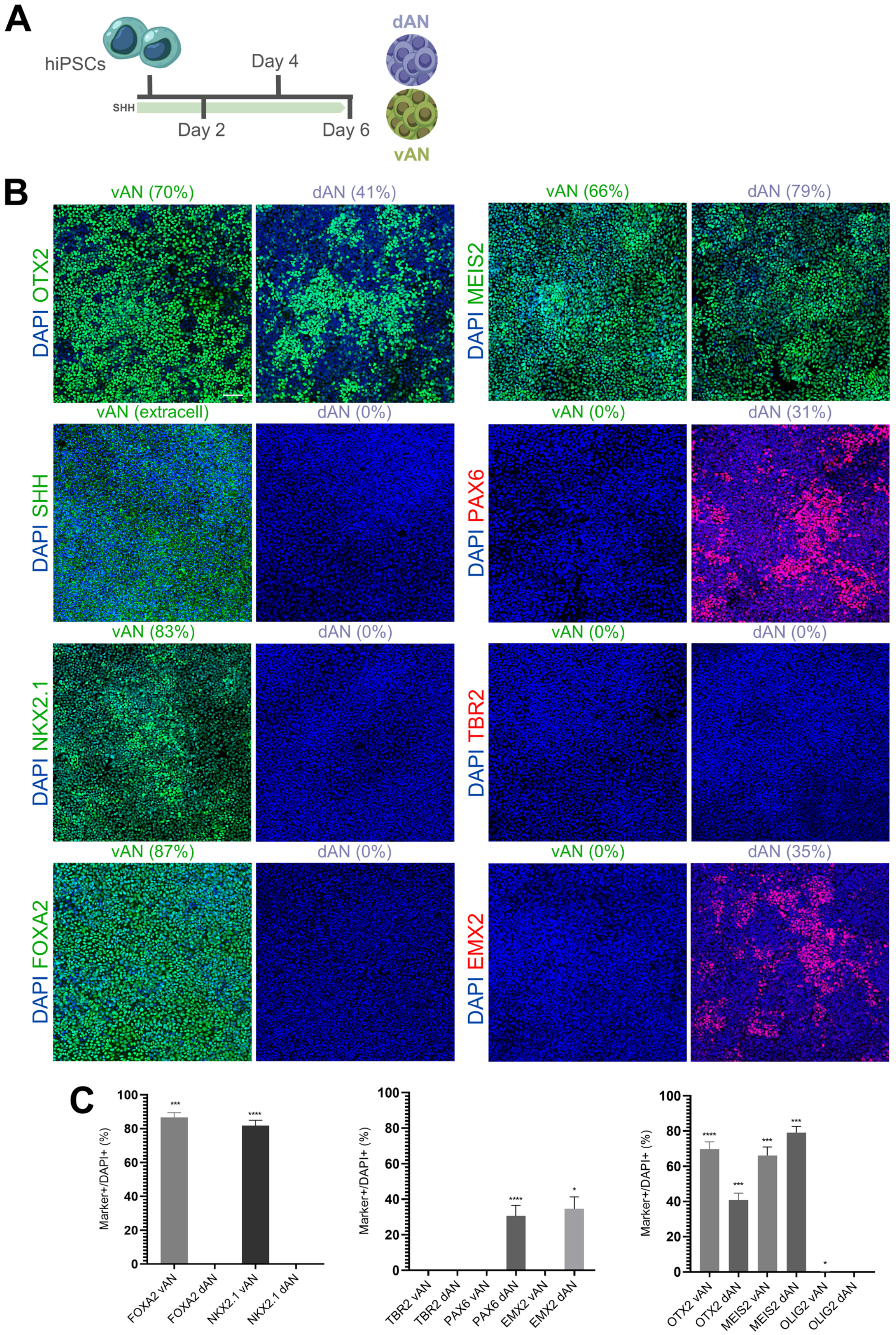
A
hiPSCs
Day 4
SHH
Day 2
Day 6
dAN
vAN
B
vAN (70%)
dAN (41%)
DAPI OTX2
vAN (66%)
dAN (79%)
DAPI MEIS2
vAN (extracell)
dAN (0%)
DAPI SHH
vAN (0%)
dAN (31%)
DAPI PAX6
vAN (83%)
dAN (0%)
DAPI NKX2.1
vAN (0%)
dAN (0%)
DAPI TBR2
vAN (87%)
dAN (0%)
DAPI FOXA2
vAN (0%)
dAN (35%)
DAPI EMX2
C
Marker+/DAPI+ (%)
FOXA2 vAN
FOXA2 dAN
NKX2.1 vAN
NKX2.1 dAN
TBR2 vAN
TBR2 dAN
PAX6 vAN
PAX6 dAN
EMX2 vAN
EMX2 dAN
OTX2 vAN
OTX2 dAN
MEIS2 vAN
MEIS2 dAN
OLIG2 vAN
OLIG2 dAN

**Figure S3: Assessment of cell identity at the mid-point of the differentiation protocol (day 6) in LON line.**

**(A)** Schematic representation of the differentiation protocol for generating dorsal (dAN) and ventral (vAN) anterior neuroectoderm from human induced pluripotent stem cells (hiPSCs) at the halfway point (day 6). This intermediate time point provides insight into the progression of regional identity specification before the full 12-day protocol is completed.

**(B)** Immunofluorescence analysis of essential markers at day 6, confirming early dorsal and ventral lineage specification. Markers assessed include OTX2 (anterior neuroectoderm marker), MEIS2 (forebrain marker), PAX6 (dorsal forebrain marker), EOMES (early neural cortical marker), EMX2 (dorsal forebrain-specific marker), SHH (ventral marker), NKX2.1 (ventral forebrain marker), and FOXA2 (floor plate marker). Quantitative data are presented as the percentage of positively stained cells relative to total DAPI-stained nuclei, averaged across five independent images. Representative images demonstrate the distinct expression patterns between dAN and vAN conditions (scale bar: 40 μm).

**(C)** Quantitative analysis of marker-positive cells in both dAN and vAN cultures at day 6, expressed as a percentage of DAPI-positive cells. Dorsal markers (e.g., PAX6, TBR1) show significantly higher expression in dAN, while ventral markers (e.g., FOXA2, NKX2.1) are predominantly expressed in vAN. Intermediate markers, indicative of shared or transitional phenotypes, display varying levels of expression depending on the differentiation pathway. Data are presented as mean ± SEM, with statistical significance levels indicated as *$p < 0.05$, **$p < 0.01$, ***$p < 0.001$, ****$p < 0.0001$ (t-tests). These results highlight the successful induction of regional identities at this intermediate differentiation stage.

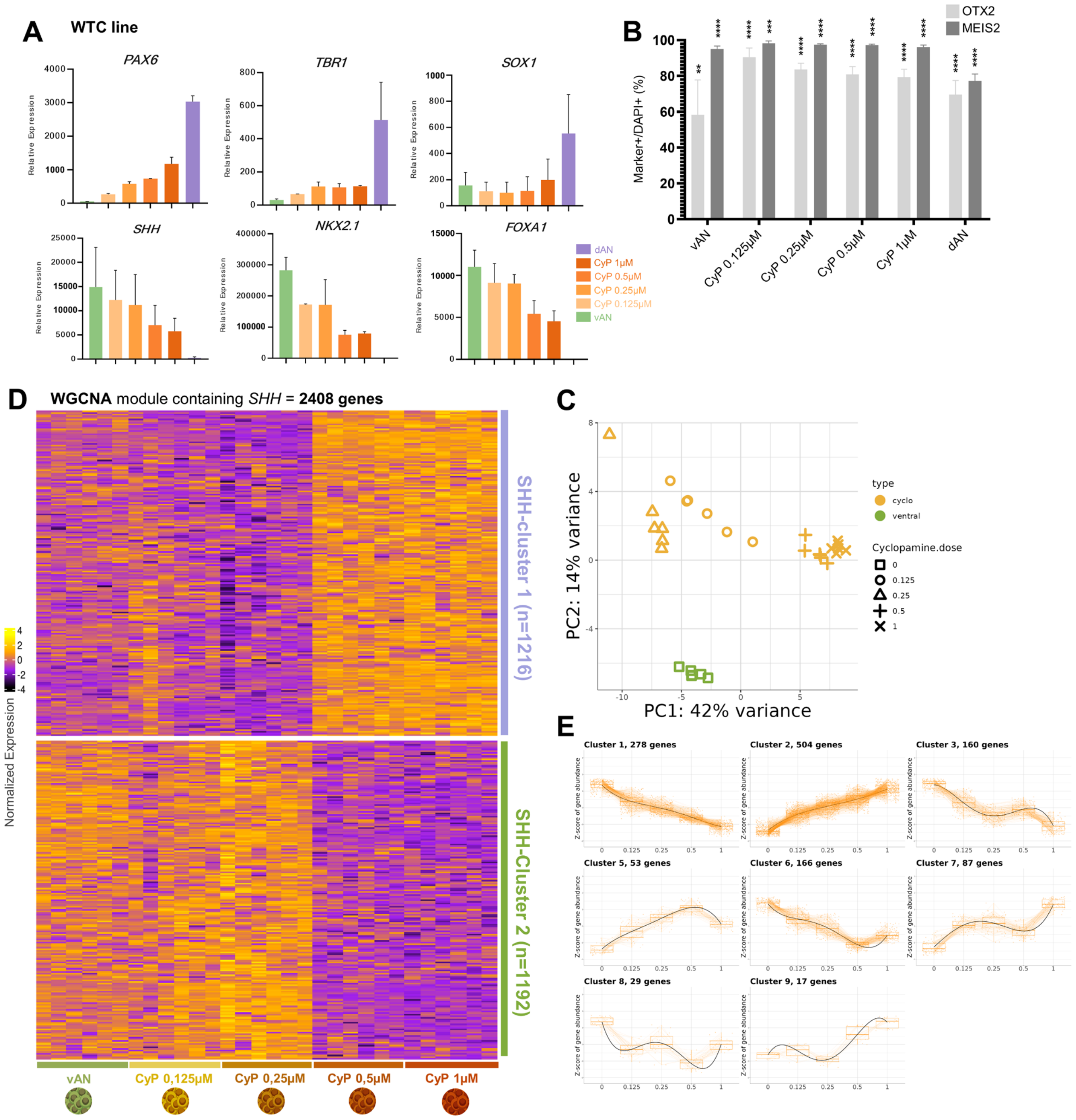

**Figure S4: Supplementary analysis of SHH pathway inhibition through Cyclopamine gradient in anterior neuroectoderm differentiation.**

**(A)** Schematic representation of the Cyclopamine (CyP) gradient protocol applied to ventral anterior neuroectoderm (vAN) cultures, with concentrations ranging from 1 µM to 0.125 µM. This gradient is used to modulate the SHH signaling pathway progressively, allowing for analysis of dorsal-ventral patterning effects in anterior neuroectoderm cells.

**(B)** RT-qPCR analysis of dorsal markers (*PAX6, SOX1, TBR1*) and ventral markers (*SHH, NKX2.1, FOXA1*) expression levels in the WTC hiPSC line differentiated under dAN, vAN, and Cyclopamine gradient conditions (1 µM, 0.5 µM, 0.25 µM, 0.125 µM) at day 12. The results illustrate a dose-dependent decrease in ventral marker expression and an increase in dorsal markers with higher Cyclopamine concentrations, indicating effective SHH pathway inhibition. Data are averaged from six replicates, with significance levels indicated as *p < 0.05, **p < 0.01, ***p < 0.001, ****p < 0.0001 (one-way ANOVA).

**(C)** Quantitative analysis of OTX2 and MEIS2 positive cells across Cyclopamine treatments and control conditions (vAN and dAN) at day 12. Values are expressed as the percentage of DAPI-positive cells, with consistent expression of OTX2 and MEIS2 indicating preserved neuroectodermal identity under Cyclopamine treatment. Statistical significance is indicated with *p < 0.05, **p < 0.01, ***p < 0.001, ****p < 0.0001. t-test.

**(D-E)** Bulk RNA-sequencing analysis at day 12 for vAN treated with gradual dosage of Cyclopamine (0, 0.125, 0.25, 0.5, and 1 μM). Cells derived from the control hiPSCs lines WTC, with five replicates.

**(D)** Principal Component Analysis (PCA) for vAN samples at day 12 treated with gradual dosage of Cyclopamine. PC1 separates sample based on Cyclopamine dosage (one-way ANOVA p.value of 7.7e-8) accounting for 42% of total variance explained.

**(E)** Heatmap showing scaled normalized expression of the genes with proper HGNC symbol within the WGCNA module containing *SHH* (2408 genes). Hierarchical clustering was applied to genes, while sample where manually organized based on Cyclopamine dosage. Clusters were defined using the cutree R function with parameter k = 2.
**(F)** Clusters from DEGpattern results for genes from *SHH*-WGCNA module on ventral and Cyclopamine samples from LON line. Each plot shows the z-score–normalized expression trajectories over time for the genes within each cluster (orange lines), with the average trend depicted in bold. Statistical significance of differences among time points is indicated by asterisks (***p < 0.001).

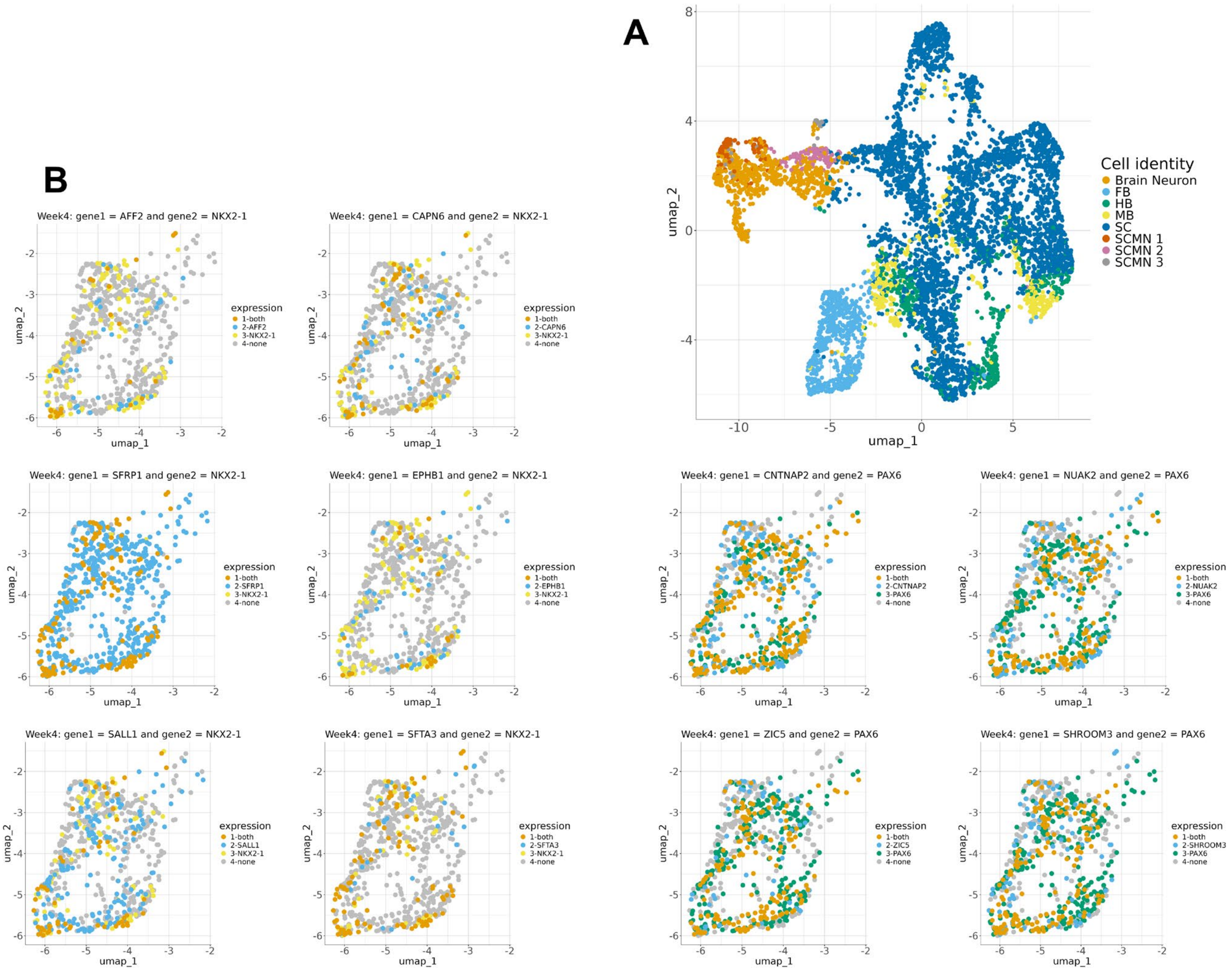


**Figure S5: Gene's selection expression verification in human brain scRNA-seq data.**

**(A)** UMAP of Zeng et al., 2023 Single cell data for Central nervous system cells at post conceptional week 4 (PCW4). Cells are colored based on cell types according to authors annotations.

**(B)** UMAP on Forebrain progenitor cells from Zeng et al. at PCW4. Cells are colored if they expressed specific genes of interest, marker genes (either *NKX2-1* or *PAX6),* or both.

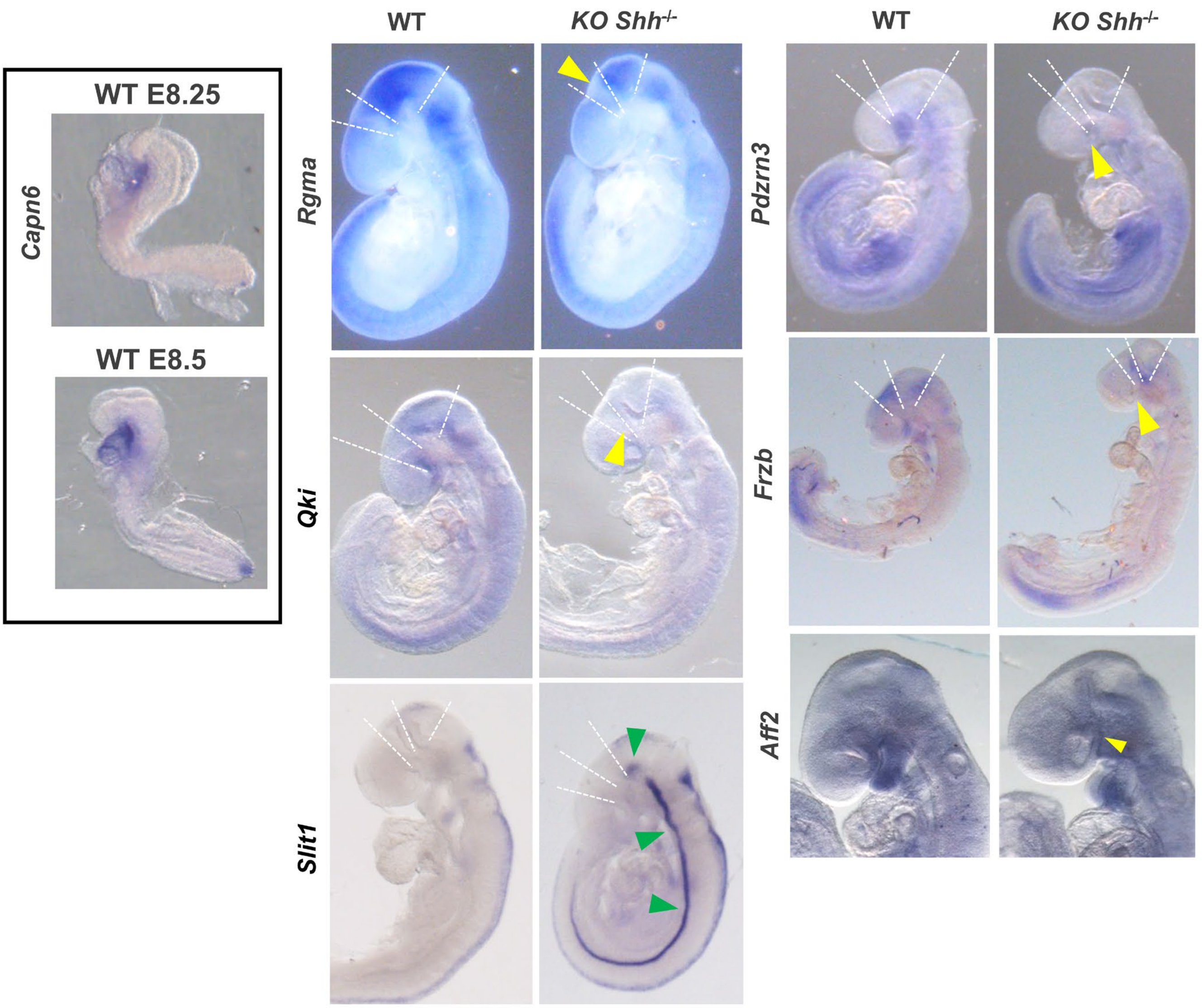


**Figure S6: Expression of several newly characterized genes in the forebrain at E9.0 and E8.25 of Wild-type (WT) and E9.0 of *Shh* mutants (*Shh* $^{-/-}$).**

The genes analyzed include *Capn6* (at E8.25 and E8.5) in WT mice; *Rgma, Qki, Slit1, Pdzrn3, Frzb, Aff2*. Black arrowheads indicate the pronounced ventral constriction at the level of the floor plate of the diencephalon and midbrain of *Shh* $^{-/-}$ embryos. Yellow arrowheads indicate the loss of mRNA expression, while green arrowheads indicate the upregulation of gene expression in *Shh* $^{-/-}$ embryos. Dashed lines represent the boundaries between telencephalon and diencephalon, the diencephalon and midbrain, and the midbrain and anterior hindbrain.
